# A Simple Way to Incorporate Target Structural Information in Molecular Generative Models

**DOI:** 10.1101/2023.02.17.529000

**Authors:** Wenyi Zhang, Kaiyue Zhang, Jing Huang

## Abstract

Deep learning generative models are now being applied in various fields including drug discovery. In this work, we propose a novel approach to include target 3D structural information in molecular generative models for structure-based drug design. The method combines a message-passing neural network model that predicts docking scores with a generative neural network model as its reward function to navigate the chemical space searching for molecules that bind favorably with a specific target. A key feature of the method is the construction of target-specific molecular sets for training, designed to overcome potential transferability issues of surrogate docking models through a two-round training process. Consequently, this enables accurate guided exploration of the chemical space without reliance on the collection of prior knowledge about active and inactive compounds for the specific target. Tests on eight target proteins showed a 100-fold increase in hit generation compared to conventional docking calculations, and the ability to generate molecules similar to approved drugs or known active ligands for specific targets without prior knowledge. This method provides a general and highly efficient solution for structure-based molecular generation.

## 1. INTRODUCTION

Computational drug design methods can be broadly categorized as ligand-based and structure-based approaches. Structure-based drug design exploits the three-dimensional (3D) information of the target protein to identify and optimize active compounds. Molecular docking is a widely used structure-based method to identify the candidate molecules that bind to a known receptor structure from a given chemical compound library containing millions of molecules. Efforts have been made to improve docking in the past three decades, including the development of empirical scoring functions, sampling algorithms, and artificial intelligence (AI)-based docking schemes^1–5^. Recent works suggested that the power of docking significantly increases with the size of the compound libraries being screened^3, 6, 7^. However, even the size of the largest compound library (a few billion) is still much smaller than the size of the chemical space which may contain about 10^60^ drug-like molecules^8, 9^.

Deep learning (DL) generative models offers a promising approach for accessing the chemical space. Molecular generative models can learn the distribution of a set of known chemical compounds and output new molecules in a highly efficient and reliable manner^10–15^. These models are essentially ligand-based, as they rely on the prior knowledge of known actives to design similar molecules that potentially bind to the same target with the possibility to fine-tune their physicochemical properties^13, 16^. To evaluate how well the new molecules bind to the target protein and prioritize them for synthesis, they can be followed up with subsequent structure-based docking calculations or molecular dynamics (MD) simulations^17^. Structure-based scoring functions were shown to be empirically better than ligand-based for molecule generation^18^. The separation of molecular generation and structure-based evaluation, however, will render the whole process inefficient and severely limits the utility of molecular generative models in practical drug discovery projects.

It’s thus tempting to develop structure-based de novo generative models that can directly leverage the 3D structure of the target protein and don’t require *a priori* knowledge of existing active compounds. One way is to perform on-the-fly structure-based evaluations and feed the results into the generative neural networks (NNs). For example, Okuno and co-workers proposed to perform docking calculations and select top compounds by the docking scores at each Monte Carlo tree search (MCTS) step in the generation process^19^. Roy and co-workers used instead a drug-target affinity (DTA) prediction model for on-the-fly evaluation^11^. The input of the DTA model is ligand-protein structural fingerprints^20^ based on docking poses and its output serves as the reward function in reinforcement learning. In another work by Roy et al, a target-specific de novo generative model is constructed by first collecting all known active compounds of the target’s homologous structures and performing docking calculations of these compounds. The docking results were then used to fine-tune a pre-trained generative model through transfer and reinforcement learning^21^. Thomas *et al*. accessed the Glide docking score to guide the REINVENT generative model. They also proposed a reinforcement learning strategy named Augmented Hill-Climb to improve the sampling efficiency of REINVENT.

Another approach for structure-based molecular generation is to generate molecules according to the geometry or shape of the target protein pocket^22–25^. Li, Pei and Lai developed DeepLigBuilder, which uses MCTS and docking scores to search and optimize both the 2D topology and 3D coordinates of molecules during generation. The coordinates were designed to be updated as the node values of MCTS such that the structure-based evaluations exempt from sampling docking poses and only involve computing the empirical scoring function^22^. Koes and co-workers constructed a conditional variational autoencoder architecture using an atomic density grid representation, which outputs ligand densities that condition on the densities of target binding sites. Molecules are then constructed from atom fitting and bond inference based on the generated ligand densities^23^. Monte Carlo tree search, on-the-fly docking or atomic density fitting is still time and resource consuming, which hinders the extensive exploration of the chemical space and thus limits the power of structure-based molecular generation. A computationally more efficient method for including the target structural information in molecular generation would represent a major advance in AI-based drug discovery.

For a specific target, docking calculations carried out on a given compound library can be considered as the process of generating labeled data for training a DL model to predict the docking scores with ligand information only. We hypothesize that such a NN model alone is sufficient to drive generative models to navigate the chemical space for extensively searching for compounds that dock favorably to the specific target. Surrogate DL docking models have been used to accurately and efficiently screen ultra-large compound libraries by training on a fraction of the library, as demonstrated by recent works including AutoQSAR/DeepChem^26^, MolPAL^27^, HASTEN^28^ and Deep Docking^29^. Whether these models are transferable in searching the entire chemical space is still unclear. Priyakumar and co-workers developed the MoleGuLAR algorithm^30^, in which predicted binding affinities were included in the multi-objective optimization for molecular generation, and the binding affinity prediction was achieved using random forest regression. Their results suggest that the prediction model lacks robustness when dealing with novel chemical substructures beyond the training dataset, and results in worse performance compared to on-the-fly docking evaluation during the generation.

In this work we demonstrate that coupling a properly trained target-specific, structure-free docking model with a molecular generative model allows effective exploration of the chemical space for discovering compounds with favorable docking scores for a specific target. It is worth mentioning that the target-specific docking models in our protocol were constructed starting with a commonly used virtual screening library, while the previous work by Roy et al. compiled and utilized for each target a compound dataset by homologous templates searching^11^. Tests on eight representative target proteins showed that our method can continuously generate diverse and novel molecules that bind well into the target pocket, although it’s important to note that in this work the binding is based solely on empirical docking calculations and not experimentally validated. The molecular generation processes were also carefully profiled. This simple method of incorporating the target structural information into molecular generative models can greatly facilitate the practical applications of structure-based molecular generations in drug discovery.

## 2. RESULTS

As illustrated in Figure 1, our protocol starts with docking calculations performed on a given compound library. In this work Autodock Vina^31^ and the ChemBridge library (743,808 compounds from its Core and Express-Pick stocks) are used. Once the docking score for each compound in the library was acquired, we trained a directed message passing neural network (D-MPNN)^32^ model to output a target-specific score which can approximate the docking score of a given molecule for this specific target protein. The D-MPNN model takes as input the molecular graph converted from the SIMLES string of the compound. Subsequently, this model is merged into a molecular generative model as the reward in the recurrent neural network (RNN) architecture. In this work we directly used the REINVENT2.0 as the underlying molecular generative model, which was trained on the ChEMBL database using maximum likelihood estimation to generate SMILES strings that corresponds to viable chemical compounds^33, 34^. Essentially the two NNs are combined for guided exploration of chemical space to extensively search for compounds with very favorable docking scores. We validated and profiled this protocol for eight representative protein targets including AmpC, D_4_, PARP1, JAK2, EGFR, PKM2, ALDH1, and MAPK1 (details in Supplementary Materials Table S1 and Fig. S1).

**Figure 1.**
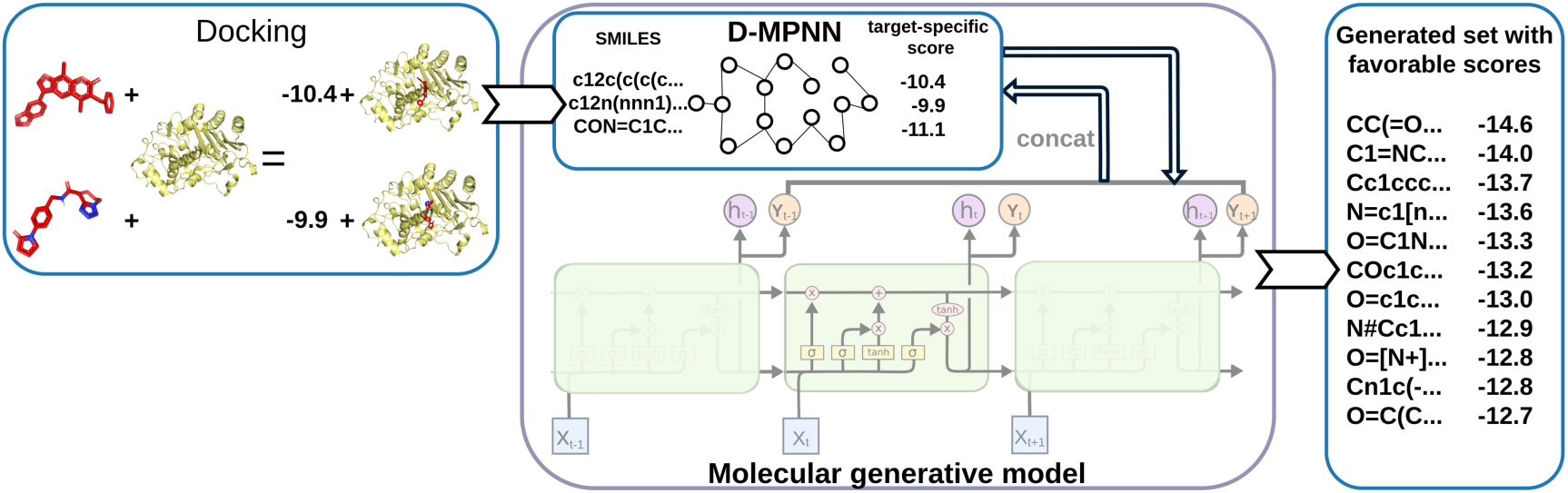
Molecular generative model guided by target-specific score. The compounds in a given compound library are docked with a specific target protein, and the SMILES and the docking score of compounds are used for training a D-MPNN model to predict target-specific scores, which is included as reward in the RNN generative model. The two NNs are thus combined to effectively explore the chemical space for searching molecules with favorable target-specific/docking scores.

### 2.1 Analysis of target-specific scoring models

For each protein, a target-specific, structure-free docking model was first trained using the docking results of all compounds in the ChemBridge library with a 9:1 split into the training and the testing sets. Such a docking model was embedded in the molecular generative model for guiding the generation of molecules that bind well to the target protein, with the protein structural information included in an implicit manner. The predictive power of the docking model should be established such that molecules with favorable target-specific scores should proportionally have favorable docking scores.

We first generated 800,000 molecules with the trained model for each of the eight targets. Reinforcement learning was conducted for 6500 epochs with a batch size of 128 until 800,000 molecules were sampled for each target. We randomly sampled 50,000 compounds from the independent ChemBridge test sets (74,381 molecules) and the generative molecule sets (800,000) to evaluate the accuracy of the target-specific scoring model. These two sets were named as C1 and V1, respectively. The target-specific scores calculated with the D-MPNN model and the docking scores computed by Vina were compared using the root mean square error (RMSE) and the Pearson correlation coefficient (PCC). As shown in Fig. 2, the target-specific and the docking scores correlated well for the C1 sets, as demonstrated by PCCs ranging from 0.78 to 0.91. The RMSEs were around 0.5 kcal/mol, except for PKM2 of which the RMSE equals 1.02 kcal/mol. In contrast, the model performance deteriorated significantly for the generated molecules (Fig. 2 and Fig. S2). Take AmpC for example, the PCC declined from 0.78 to 0.49 and RMSE increases from 0.47 to 0.79 kcal/mol comparing the C1 and the V1 sets. Similar decrease of the PCCs and nearly doubling of the RMSEs were observed for all the targets. We note that a sample size of 50,000 was large enough such that different sampling of the generative molecules leads to very similar PCC and RMSE results (Table S2).

**Figure 2.**
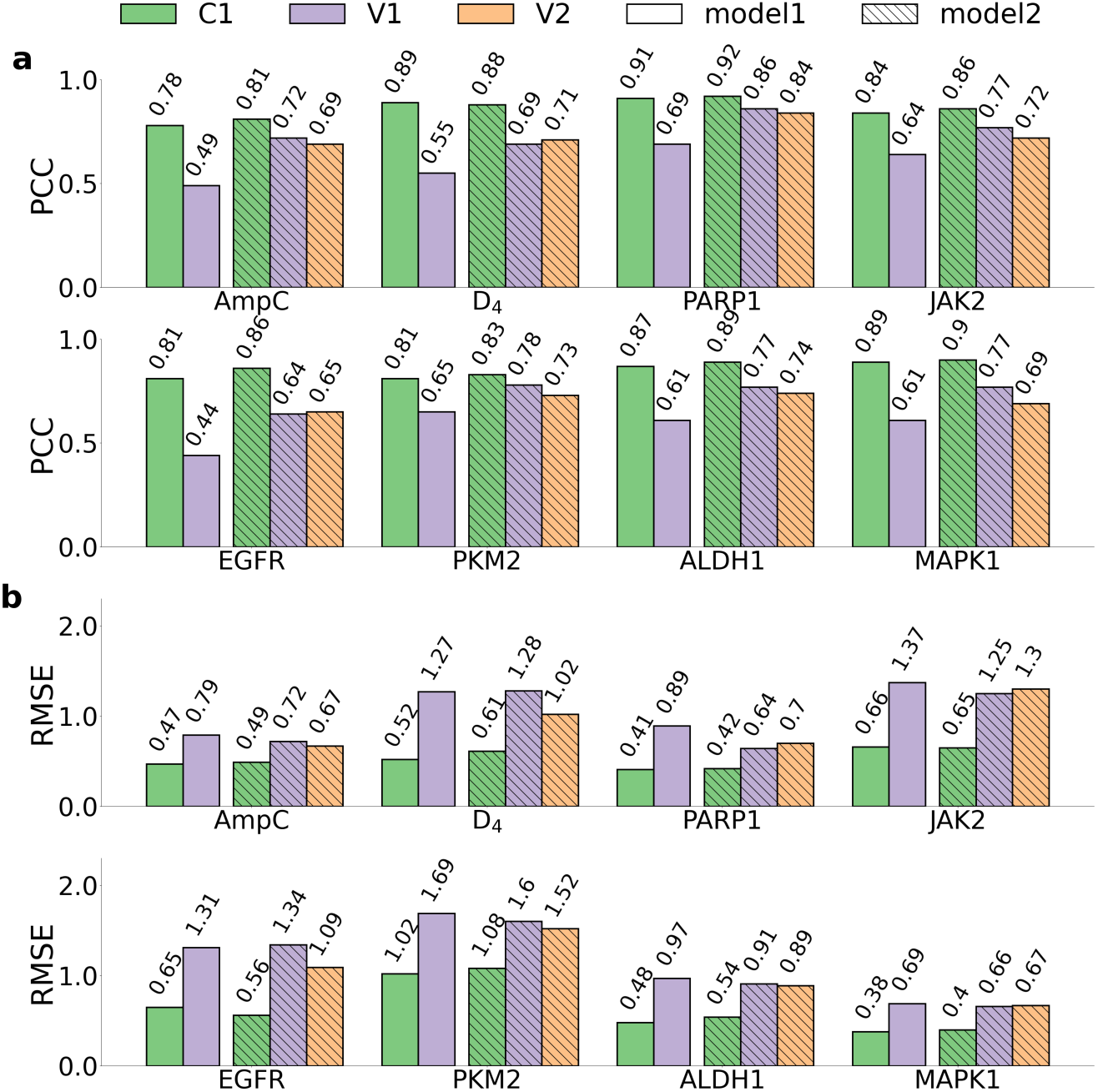
Performance of target-specific scoring models for eight target proteins. a. PCC and b. RMSE were calculated for independent testing datasets from ChemBridge, generated datasets guided by model 1 and generated datasets guided by model 2 named as C1 (green), V1 (purple) and V2 (orange), respectively. The model 1 was trained with molecules from ChemBridge library, while the model 2 was trained a mixed dataset combining ChemBridge molecules and generated molecules.

We reasoned that the decrease in the model’s ability to reproduce the docking scores was due to the fact that the V1 sets were already sampling manifolds in the chemical space that were significantly different from the starting ChemBridge library. While it is in general difficult to visualize manifolds in the chemical space, a 2D principal component analysis (PCA) projection using the molecular quantum numbers (MQNs)^9, 35^ illustrated the generated V1 sets did shift individually away from the ChemBridge library (Fig. S3). This can also be partially inferred from the difference in docking scores. As shown in Fig. S4, the distribution of the Vina docking scores for the V1 sets already shifted towards more negative values for all the eight targets by 0.6 to 2.5 kcal/mol. It’s interesting to observe that generated molecules were slightly larger and more lipophilic than compounds in the starting ChemBridge library, as reflected in the distributions of their molecular weights (MWs) and computed logP^36^ (Figure S5). We note that it’s well known that larger molecules in general have more negative docking scores, thus we restrained the MW at 650 Da to prevent generating very large molecules.

A straightforward resolution would be to re-train the target-specific scoring model using an extended set of molecules that covers better the target-specific manifolds in the chemical space. Specifically, for each target 500,000 generated molecules and 500,000 molecules in the ChemBridge library were randomly selected and mixed for a second round of training using the same D-MPNN architecture and hyperparameters. The re-trained models (model 2 in Fig. 2) were used for guided exploration of the chemical space and generated the same number of molecules. Similarly, we sub-sampled 50,000 out of the 800,000 generated molecules as the generative set V2, which were used together with C1 and V1 to evaluate the performance of the new target-specific scoring models. As illustrated in Fig. 2, the new models improved slightly on the original C1 set in terms of PCCs. Significant increase in the PCCs and decrease in the RMSEs were observed for the V1 sets, which can be understood as the training data now included more molecules that were likely to bind to the specific target. We note that the calculations of PCCs and RMSEs for the V1 sets with the model 2 excluded any molecule used in model training. Most importantly, the performance on the V2 test set was comparable with the V1 generated set predicted by the re-trained model 2. The PCCs range from 0.65 (EGFR) to 0.84 (PARP1) and the RMSEs vary from 0.66 (MAPK1) to 1.5 kcal/mol (PKM2) for the V2 sets. The model performance was still relatively worse for generated molecules compared with the ChemBridge library, which hints on the higher intrinsic complexity in the generative sets that might impose more difficulties for NNs to learn.

### 2.2 Generation guided by target structural information yields molecules with notably more favorable docking scores

A NN model with reasonable correlation between target-specific and docking scores allows to sample the chemical space efficiently and accurately. To benchmark the binding affinity of newly generated molecules comparing with compounds in the ChemBridge library, we selected equal number of molecules (743,808) sequentially from the molecular generation guided by target-specific scoring model. Figure 3 compared the numbers of molecules with Vina docking score above a range of thresholds in the generated sets and the ChemBridge library. For all eight target proteins, molecular generation results in molecules with significantly better binding affinities as evaluated by docking calculations. For targets such as D_4_ and PARP1, molecules with docking score as negative as -17 kcal/mol were identified.

**Figure 3.**
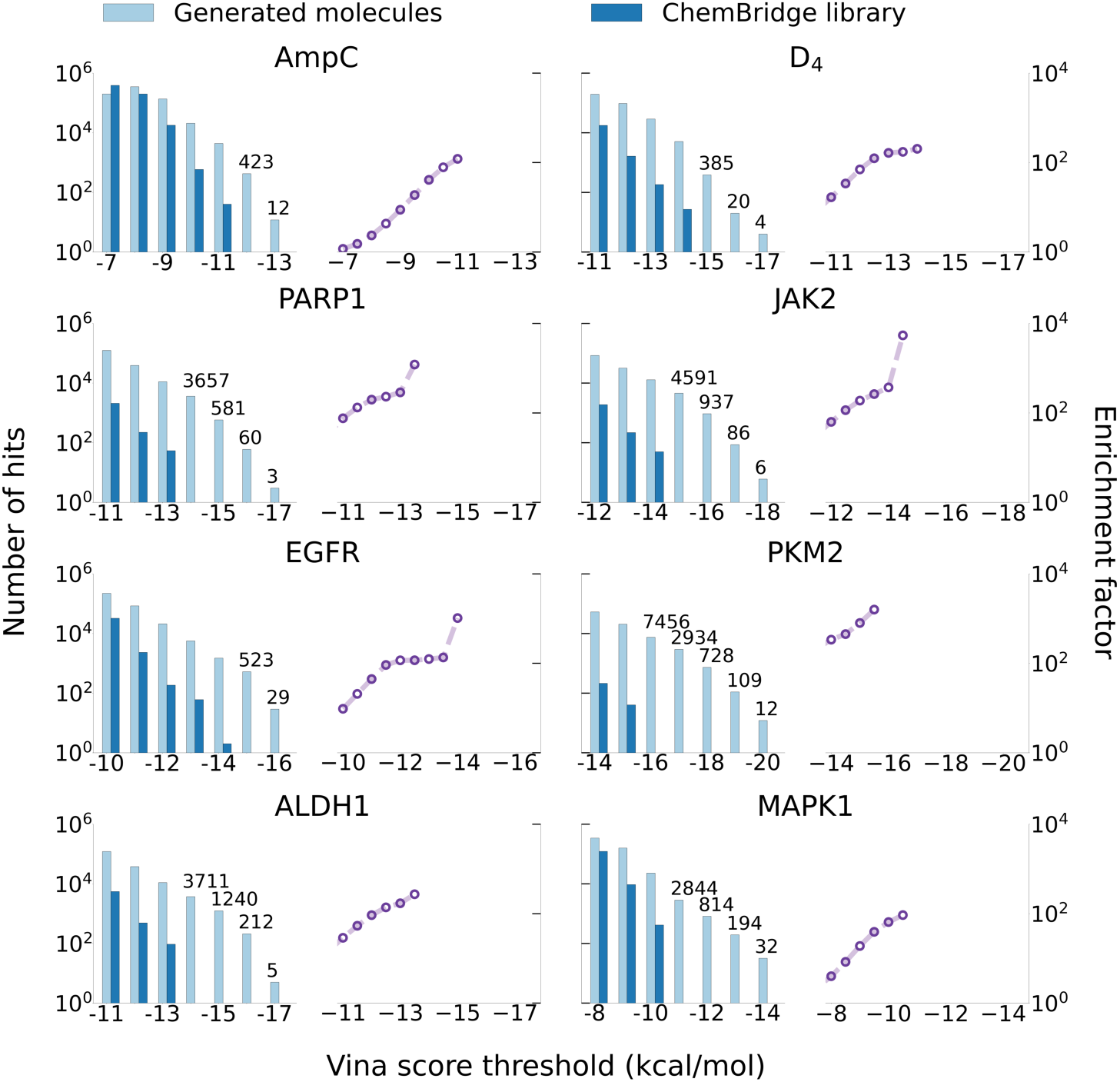
Comparison of generated molecules (light blue) and ChemBridge compounds (blue) with respect to their docking scores. The numbers of hits identified using different score thresholds were compared for the eight target proteins. Enrichment factors were plotted as the purple lines.

To quantify the enrichment of generated molecules against the ChemBridge library, enrichment factors were shown in right panels of Fig. 3 which are defined as the ratios of the number of candidate hits above a given docking score threshold. For example, at the threshold of -13 kcal/mol, our protocol yielded 164, 287, 188, 124, 116 and 170 times more potential hits compared with ChemBridge for D_4_, PARP1, JAK2, EGFR, PKM2 and ALDH1. Using the same threshold for AmpC and MAPK1, there is no candidate hit with ChemBridge docking calculations but 12 and 226 potential hits from molecular generation. If we set the threshold to the docking score of the 100th ranked ChemBridge hits, the enrichment factor ranges from 88 (AmpC) to 488 (PKM2). The increase of two orders of magnitude illustrated the power of molecular generation guided by target-specific scoring in generating the promising compounds that bind with a specific target protein.

We further decomposed the Vina docking scores into hydrogen bond, hydrophobic interaction and steric interaction terms^31^, and analyzed how the generation protocol realized favorable docking. Figure 4 compared the distribution of three energetic terms of the top 1,000 molecules from the generation and the ChemBridge library. In general, all three terms were shifted to lower (better) values for generated molecules, suggesting that our protocol exploited all possible ways to generate molecules that can fit better into the target pocket. The values of steric interactions were lowered by 1.3-4.0 kcal/mol for all targets except MAPK1 with a minor decrease of 0.2 kcal/mol. It’s interesting to note that for some targets (D_4_, PARP1, JAK2, and ALDH1) the generated molecules had stronger hydrophobic interactions, but very similar hydrogen bond interactions compared to the ChemBridge hits. Meanwhile, for some targets (AmpC and MAPK1) molecules generated by our protocol can form more hydrogen bonds but failed to establish more hydrophobic interactions with the target proteins. Detailed inspection on their pockets shows that the pockets of D_4_, PARP1, JAK2 and ALDH1 are very hydrophobic while those of AmpC and MAPK1 contain more polar residues (Fig. S1). For EGFR and PKM2, the generative models were able to establish both more hydrogen bonds and stronger hydrophobic interactions. It’s encouraging that the protocol learns the pocket environment for guided generation of molecules that fit specifically for the target. The generation process indeed feels the shape and interaction pattern of the target pocket, although the 3D structural information is introduced in an implicit way.

**Figure 4.**
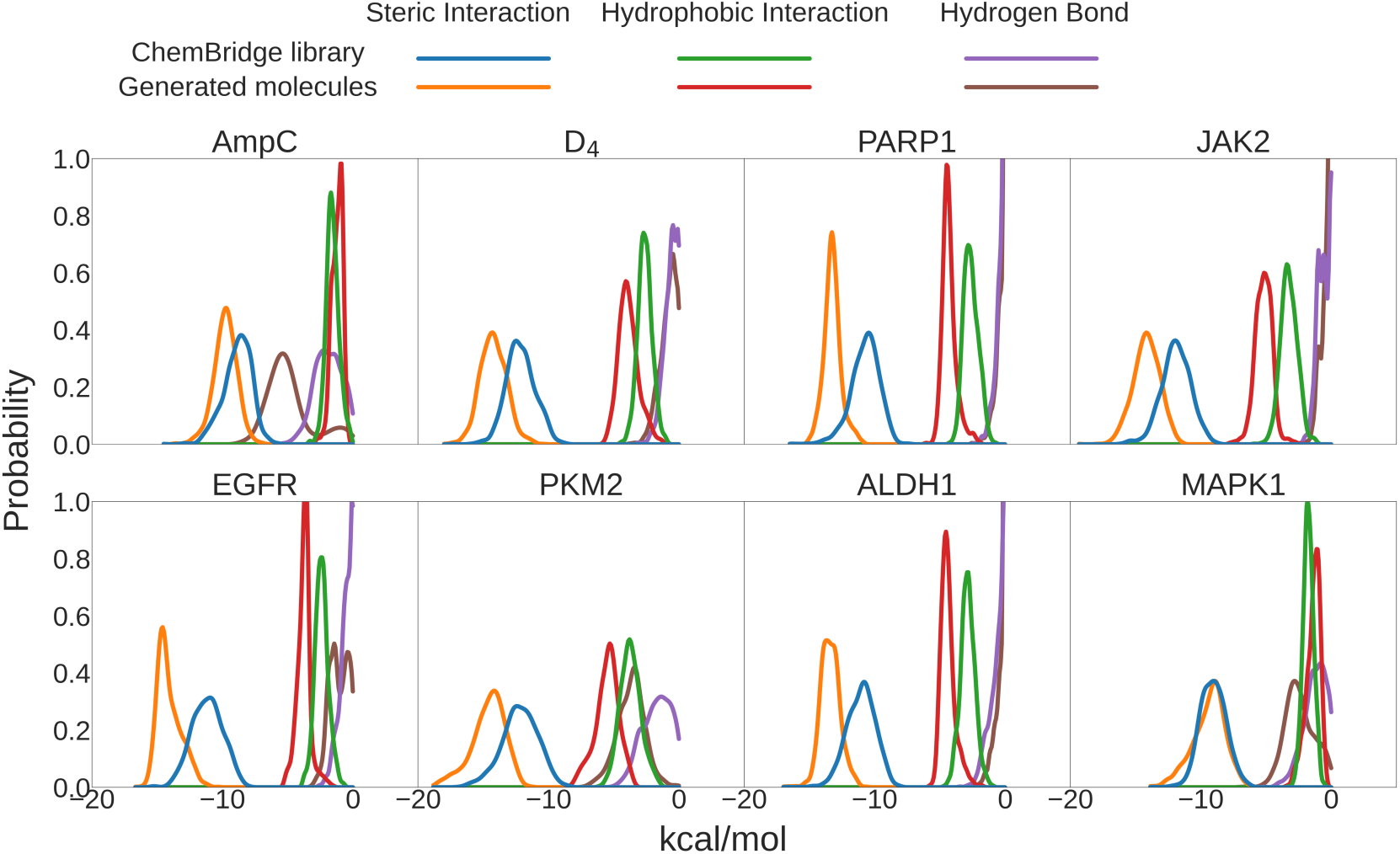
Comparison of Vina score components for generated molecules and ChemBridge compounds. The Vina docking scores were decomposed into steric interaction (blue and orange), hydrogen bond (green and red) and hydrophobic interaction (purple and brown).

We note that for each target protein, the scoring model was trained using the docking results of a combined total of 1,243,808 molecules (743,808 from ChemBridge and 500,000 from the V1 set). Conducting docking at this scale is a common practice in drug design, owing to its inherently parallelizable nature. In this work, the docking calculations required about 33 hours using 2000 CPU cores (Table S3). Once the target-specific scoring model was obtained, the structure-based molecular generation can be quite efficient, generating 800,000 molecules within a few hours (Table S3). An additional advantage is that the trained D-MPNN model can be re-used for multiple reinforcement learning runs.

One of the target studied here, JAK2, was also used in a previous study^21^ to demonstrate that molecules can be generated with favorable Vina docking scores. Applying the same drug-like physicochemical filters with the ref^21^ to the generated molecules in this work leads to 524,282 molecules. Figure S6 compares the distribution of Vina docking scores for these 524,282 molecules and the 6,106 generated molecules reported in ref^21^. The first quartiles equal -11.3 and -10.1 kcal/mol, while the medians equal -10.5 and -9.3 kcal/mol, respectively. Note that to eliminate the possible impact from different settings in Vina calculations, we recomputed the Vina scores for reported molecules^21^ and obtained consistent numbers (Fig. S7). The ability to discover molecules with more favorable docking scores is mainly due to the significantly more extensive searching in the chemical space guided by an accurate surrogate model, as evidenced by a steady increase in the average docking scores for generated molecules during the first 1000 epochs, which corresponds to 128,000 molecules (Fig. S8). In general the docking scores of generated molecules by our simple protocol are significantly more negative than commonly reported generative models^25, 37^. We note that a Vina docking score of -17 or -18 kcal/mol is rarely seen in the literature and we shall discuss it later.

### 2.3 Most generated molecules are novel chemical compounds

We next evaluate the novelty and diversity of molecules generated with the guidance of target-specific scoring model. The similarity of the top 100,000 generated molecules with all the compounds in the ChEMBL database^38^, the ChemBridge library and the ZINC20 database^39^ were assessed using the Tanimoto coefficient (TC) with ECFP4 fingerprint. For each generated molecule, we computed the TC similarity against each curate molecules in each database and reported the maximum TC values in Fig. 5. We note that ChEMBL was used to train the REINVENT2.0, which is used as the baseline generative model in this study. The ChemBridge library was used for the preliminary training of target-specific scoring models, while the ZINC20 database (1.6 billion molecules) represents the largest purchasable collection of chemical compounds.

**Figure 5.**
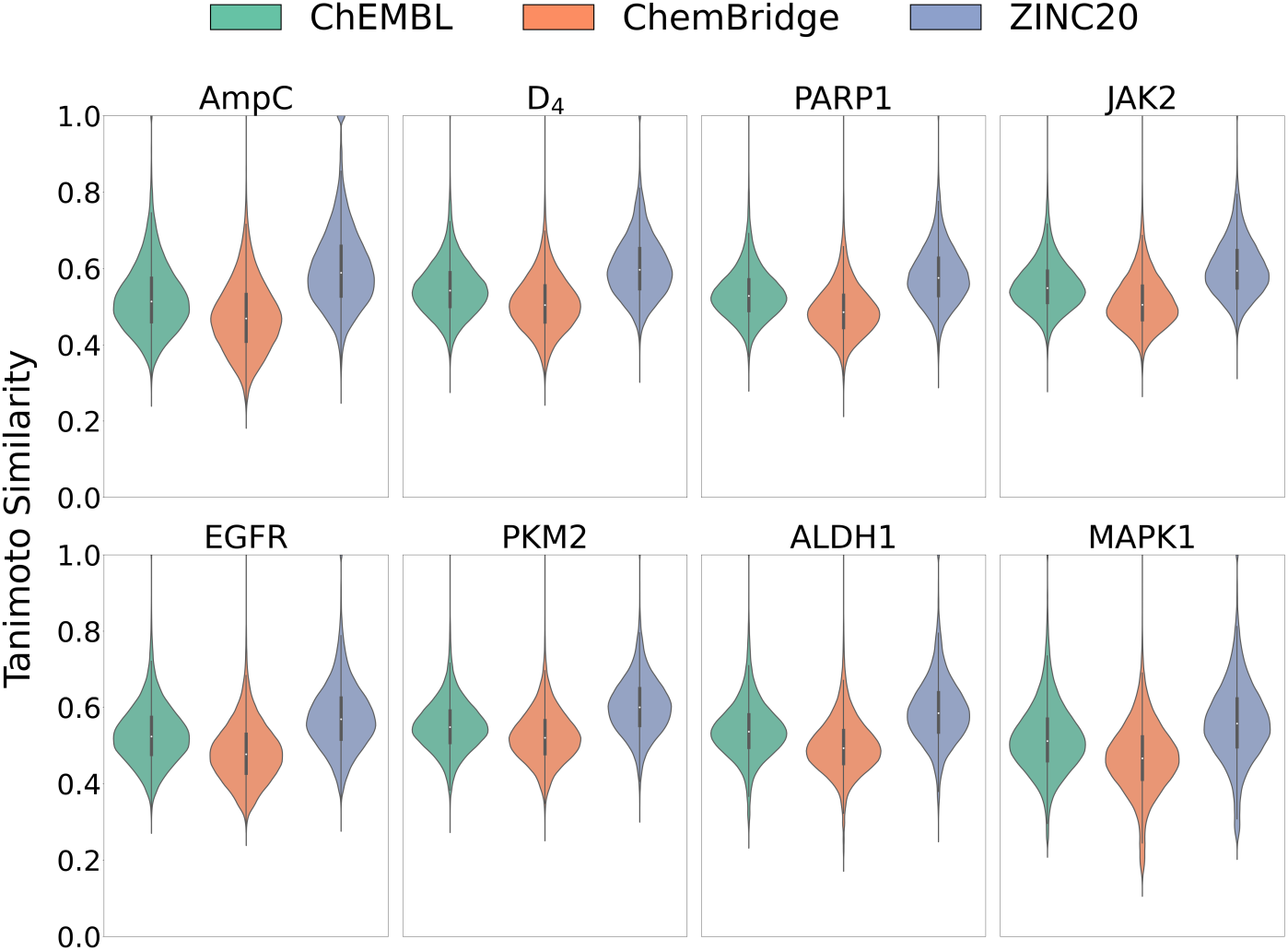
Distribution of the Tanimoto coefficient similarities between the top 100,000 generated molecules with ChEMBL (green), ChemBridge (orange) and ZINC20 (purple) libraries which included about 2.2 million, 0.7 million and 1.6 billion compounds, respectively. Each individual violin plot represented the largest TC values comparing the generated molecules with compounds in the three databases for the eight target proteins, respectively.

As shown in Fig. 5, the median values of the maximum TC similarity for the eight targets range from 0.51 to 0.55, 0.47 to 0.52, and 0.56 to 0.60 when compared with ChEMBL, ChemBridge and ZINC20, respectively. Considering the substantial quantity of molecules in these databases, such TC values indicate that most generated molecules are dissimilar with compounds in public databases and represent new molecular entities. In general, the similarities with ZINC20 compounds were higher, which is reasonable due to the significantly larger size of the ZINC20 database. There were few molecules with TC=1, indicating that these molecules generated from scratch happen to be exactly the same as some curated in the databases. In particular, 1262, 275, 124, 130, 286, 127, 318 and 324 generated molecules were found to be identical to one of the ZINC20 compounds for AmpC, D4, PARP1, JAK2, EGFR, PKM2, ALDH1 and MAPK1, respectively (Table S4). Synthesizing novel molecules is in general considered to be a bottleneck for generative models in drug discovery, and the possibility to generate purchasable compounds can be advantageous in the early discovery phase to test the computational models with the costs and uncertainties in synthesis minimized. Also presented in Table S4 are the number of identified hits that are present in the ChemBrdige and ChEMBL databases, which were used for training but could also be considered as subsets of ZINC20. On average, about 15 hits were recovered in ChemBridge, representing approximately 0.002% of the docking data used in training the target-specific scoring models.

Docking calculations showed that these generated purchasable compounds had favorable docking scores and poses. As typically dozens of compounds would be purchased and tested for bioactivity, we summarized in Table S4 the Vina docking scores for top 20 and top 50 purchasable hits and their values were less than -10 kcal/mol for all targets. Docking ultra-large libraries such as ZINC is notoriously difficult,^3^ and our protocol might serve as a detour to tackle this challenge. However, it should be noted that the effectiveness and efficiency of current protocol in identifying top-scoring molecules require further validation.

### 2.4 Comparison with known active compounds

The Tanimoto similarity calculation with the ChEMBL database allows further profiling of generated molecules as ChEMBL contains bioactivity data. Experimentally reported active ligands were collected from the ChEMBL database for all eight target proteins, and similar compounds can be identified from the top 100,000 generated molecules except for the AmpC target (Table S5). Figure 6 illustrates for each target a representative case by comparing the Vina docking poses of a known active compound and a similar generated molecule. Minor differences were observed, such as the para-chlorination of a phenyl group in ChEMBL1502164 for MAPK1, or the replacement of the cyclopropyl group with a cyclobutyl group in ChEMBL521686 for PARP1. Most importantly, very high ligand 3D similarities, as measured by LS-align^40^ alignment, were found. This suggests that the select generated molecules had interactions with the target pockets that were sufficiently similar to those of their nanomolar inhibitors, as shown in Fig. 6. These molecules in fact had more negative Vina docking scores, and it’s worth pointing out that no information on active ligands was used during the entire molecular generation process.

**Figure 6.**
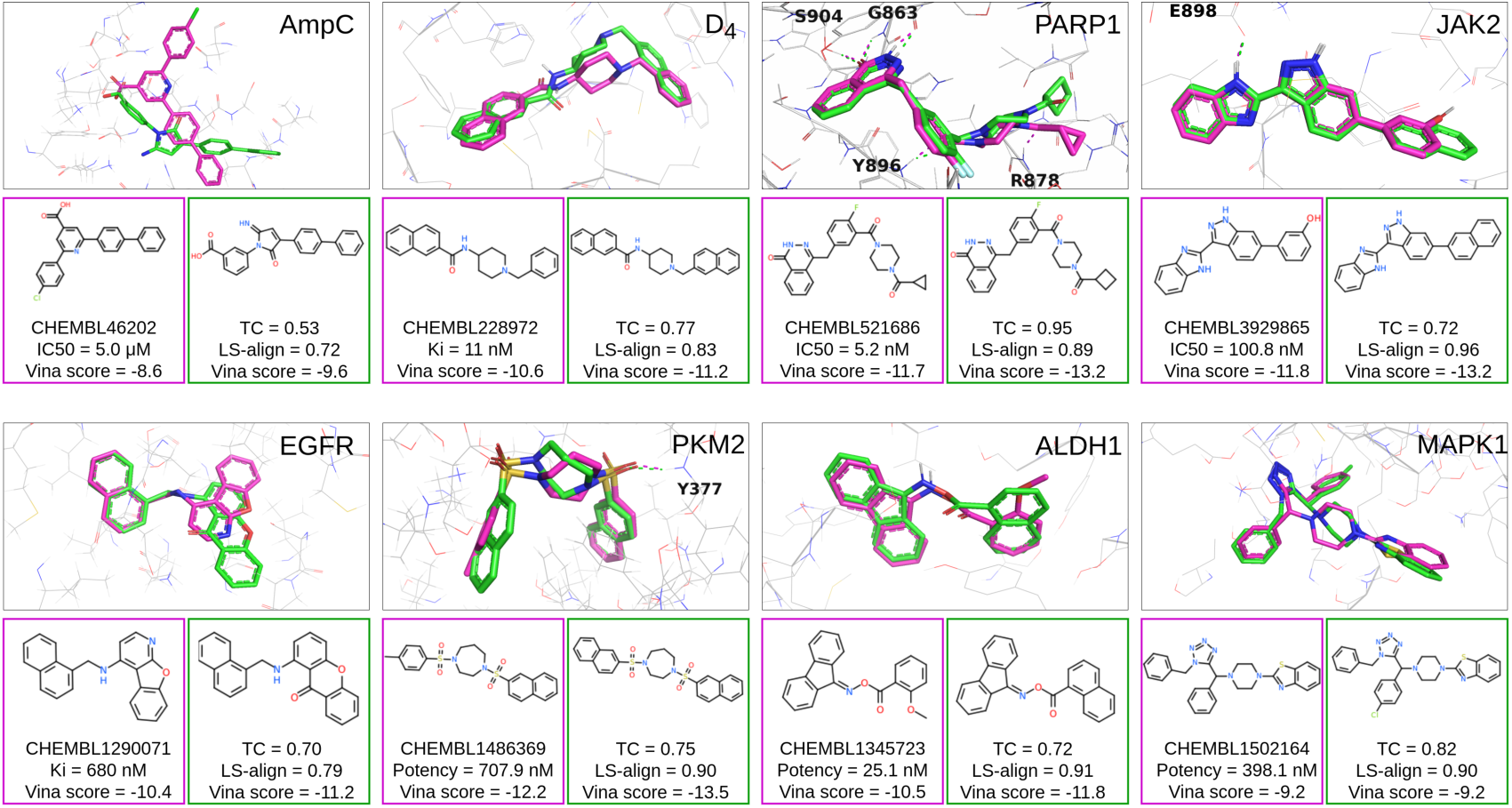
Comparison between generated molecules (green) and known active compounds (magenta) for eight target proteins. The docking poses were generated by Vina, the 2D similarities represented by TC, and the 3D similarities calculated with LS-align rigid body alignment.

One case in point, the PARP1 inhibitor ChEMBL521686, is a marketed drug (Olaparib) for treating BRCA-mutated cancers. We note that 53 out of the top 100,000 PARP1-specific generated molecules had a TC of 1 when compared to ChEMBL compounds (Fig. 5), and 21 of them (about 39.6%) were found to be identical to previously reported inhibitors for the PARP1 target protein (shown in Fig. S9). These generated molecules shared a similar phthalazin-1(2H)-one scaffold while differed in the chemical moieties to establish interactions with key residues such as Y896 and R878 (Fig. 6). This result illustrated that the generation protocol is capable of performing local search on molecular substructures. Combining with customized filters, target-specific generative models can be used to explore structure-activity relationship in drug discovery projects.

We note that ChEMBL521686 from the ChEMBL dataset was used for training the baseline generative model (REINVENT2.0), and it is not in the ChemBridge dataset used for training the D-MPNN model. The training of REINVENT utilizes the chemical information of compounds in the ChEMBL library; however, it does not leverage information on whether a compound is active or inactive for a given target. Therefore, no prior information on the activity of ChEMBL521686 with respect to PARP1 was included in the generation protocol. To further illustrate the importance of implicit structure-based affinity evaluation, we carried out five independent runs of REINVENT2.0, each generating 100,000 molecules, and compared the results in Table S6. It was observed that these baseline generations resulted in a considerably larger number of generated ChEMBL molecules, and the percentage of recovering ChEMBL hits for the PARP1 target dropped to a mere 0.2%. If the molecular generation was not guided by the target-specific scoring model, the recovered hits generally exhibited less potency in terms of their experimental affinities (Fig. S10).

### 2.5 How many scaffolds can be sampled for a given target?

The high level of efficiency of such a generation protocol opens the possibility to address this question by continuously cranking the generative model. Using PKM2 and ALDH1 as examples, we profiled the ability of our generation protocol to continuously explore the chemical space for potential hits with unique scaffolds for a given target. Figure 7 shows the number of unique potential hits that can be discovered every GPU hour on a single NVIDIA A40 card. A variation of Murcko scaffold^41^ was adopted by converting all heavy atoms into sp3 carbons, and a generated molecule was defined as unique if its Murcko scaffold was not present in the previous epochs during the generation. Furthermore, it was considered to be a potential hit if its target-specific score was better than the 10^th^ percentile of the first 800,000 generated molecules as analyzed in previous sections, which equals -13.9 and -11.3 kcal/mol for PKM2 and ALDH1, respectively.

**Figure 7.**
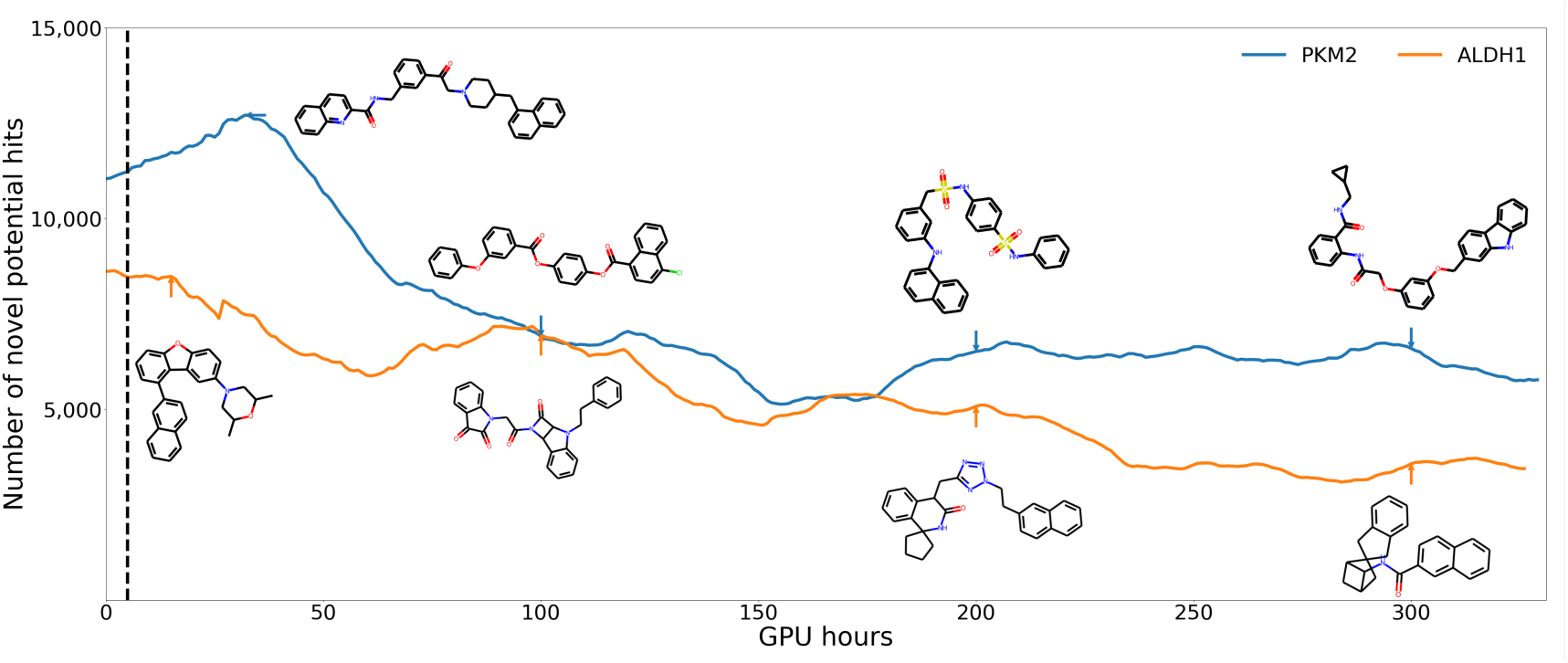
The number of potential hits with unique scaffolds generated for PKM2 (blue) and ALDH1 (orange) every GPU hour. Representative molecules were plotted and the dash line represents the runtime for generating 800,000 molecules (about 5 GPU hours).

We note that the molecular generation processes already incorporated penalization for repeating scaffolds (see Methods for details). This is also the reason that GPU hours instead of generation epochs were used in Fig. 7 as the runtime for each epoch gradually decreased. To generate 800,000 molecules, typically 6,500 epochs were employed which took about 5 hours (dash line in Fig. 7), and about 60,000 unique potential hits were identified based on our criteria. When we kept running the generative models, potential hits with unique scaffolds continuously emerged, although with a gradually declining pace. Peaks were found after 32 and 15 GPU hours for PKM2 and ALDH1, with the first unique scaffolds at the peaks marked by blue and orange arrows, respectively. Also listed were the first unique scaffolds after 100, 200, and 300 GPU hours. Over a runtime of 300 GPU hours, millions of potential hits with unique scaffolds can be generated. This also demonstrates the efficiency of the guided structure-based molecular generation once the surrogate docking model is trained. It’s inspiring that comprehensive exploration of the chemical space can yield in a significant number of molecules that fit well into a given pocket, which implies a vast margin for small molecule drug design. We do observe that the molecules generated later exhibit more complicated stereochemistry, including spiro and bridged compounds.

Finally, we would like to emphasize that the activity of a compound for a given target was evaluated solely based on docking calculations in this work, without any experimental validation. A favorable docking score does not necessarily imply strong binding affinity, and may often not result in the expected bioactivities. However, docking has been a cornerstone in drug discovery for decades. Recent work by Lyu and co-workers^3^ demonstrated that the experimental hit rates were monotonically related to docking scores when performing large-scale screenings, and highlighted the importance of accessing expansive chemical spaces through docking with ultra-large compound libraries. The computational protocol proposed in this work can thus also be viewed a novel way to use docking in rational drug design.

## 3. CONCLUSION AND DISCUSSIONS

In this work, we proposed a simple way to incorporate target 3D structural information into molecular generative models. The basic idea is to use docking calculations to train a target-specific scoring model, which is then combined with a molecular generative model to search in the chemical space for compounds that bind favorably to the target. The key component of this protocol is to construct for each target a dataset that includes plenty of diverse molecules that have the potential to bind favorably, and to use this dataset for training the D-MPNN scoring model. To this end, both the docking scores and the molecules used for the training are target-specific. This target-specific dataset itself can be obtained by preliminary training and generation starting from a common public compound library. Our results, using eight protein targets as examples, demonstrate that the algorithm allows guided exploration in the chemical space with high level of efficiency. While the idea of coupling a generative model with a predictive model is not new^15^, in this work we illustrated that simple training of a surrogate DL docking model suffers from poor transferability when combined with a generative model for searching bioactive manifolds in the chemical space for a given target. We provide a solution to improve the transferability, which would further unleash the power of generative models in drug discovery. Unlike with other structure-based generations, our model can efficiently generate millions to billions of molecules according to the pocket environment of target protein without relying on collecting the active and inactive for training the classification model. Our model entirely accomplishes the target-specific design by two combined NN models to learning the interaction between target proteins and respective compounds.

The connection and difference with DL-based docking methods^26–29^ shall be discussed. The aim of these methods is to reduce the computational cost of docking calculations in screening a given ultra-large compound library. Docking results on a small fraction of the library are used to build a structure-free DL docking model for predicting docking scores with the 1D or 2D representations of ligands, which leaves out the processes of sampling docking poses and computing interaction energies. The D-MPNN architecture in this work is based on that of MolPAL^27^. The transferability issue was also discovered by the DL docking methods such that active learning is found to be essential for them to work^27, 29^. Another example is V-Dock^42^, which uses a surrogate docking model as part of the linear objective function for evaluating molecules generated and optimized by the conformational space annealing algorithm^43^. Significant decrease in the correlation between predicted and actual docking scores was observed between training set and generated molecules^42^. Our solution doesn’t involve active learning, although two rounds of training were employed. We consider the 1st round of training and the molecular generation with this model a preliminary step for the creation of a dataset that represents molecules that might dock favorably with the specific target. By using predictive docking model in the context of molecular generation, we can directly explore the bioactive manifolds, which are continuous and structured in the high-dimensional chemical space. This contrast scoring a curated compound library, which corresponds to sporadic points in the space.

We used Autodock Vina in this work and reported that molecules can be generated with very favorable docking scores such as -18 or -20 kcal/mol. Docking scores are empirically developed using experimental protein-ligand complex structures and binding affinities, and it’s well known that they can deviate - sometime significantly - from experimental affinities^44^. Manual inspection on generated molecules with extremely favorable Vina scores indeed find some of them apparently won’t bind. Figure S11 listed examples of generated molecules for ALDH1 with Vina scores ranging from -11 to -17 kcal/mol. To obtain very negative docking scores, molecules diminished in features and became aromatic hydrocarbons, which presumably maximize the number of ligand-protein contacts for the long-range steric attraction (Eq. 7 in Ref. 32) and the hydrophobic interactions, and minimize the number of rotational bonds (Eq. 9 in Ref. 32) ^31, 45, 46^. This demonstrates that the generation protocol presented in this work are powerful enough to exploit the empirical nature of docking scores, while also indicates the need for further evaluation or filtering of the generated molecules for practical use.

Our protocol is a universal workflow for guiding molecular generative models with structure-based binding calculations, and molecular docking approaches other than Autodock Vina can be easily adopted. To illustrate this, we substituted Vina with DOCK6 or the Vinardo docking score and tested the protocol on the EGFR target (Figure S12 and S13). Similarly, we observed that a second round of training, incorporating the docking results of generated molecules as training data, led to improvement in the predictive power of the target-specific D-MPNN models. Using the docking scores of the 100th ranked ChemBridge hits as the threshold, the enrichment factors were 116, 608 and 1,116 for the target-specific molecular generation employing Vina, Vinardo, and DOCK6, respectively. Notably, the DOCK6-derived model showed the highest correlation between target-specific and docking scores, with a PCC of 0.81 (Fig. S12). It’s important to reiterate that, in this work, the activity of a compound was evaluated solely with the docking score. Therefore, performance in terms of experimental affinity will depend critically on the quality of the docking model used, which can vary significantly across different targets.

To increase accuracy, the protocol also allows for the replacement of docking with more sophisticated absolute or relative free energy perturbation (FEP) calculations to infer the binding affinity of a compound to a specific target protein^47–49^. However, the predictive power of FEP calculations might be constrained by limited molecular diversity as one typically enumerates restricted sets of R-groups^49^. To alleviate the associated computational cost one can fine-tune the last layers of “model 2” with FEP results, and the same strategies can also be used to integrate experimental affinity data. Similarly, the protocol is versatile for using alternative molecular generative models beyond REINVENT, with the possibility of “focused” generation with specific core structures, and using beam search score^50^ or perplexity score^51^ to rank the generated molecules according to the probability of their SMILES tokens. In conclusion, we introduce a general and efficient new method for structure-based molecular generation.

### Data and Code Availability

All source codes are available at https://github.com/JingHuangLab/SWIT, together with data files to reproduce the results in this manuscript. The ZINC20, ChEMBL and ChemBridge datasets were obtained from https://zinc20.docking.org/tranches/home, https://chembl.gitbook.io/chembl-interface-documentation/downloads, and https://chembridge.com/screening-compounds/lead-like-drug-like-compounds, respectively.

### Supporting Information Available

Protein-ligand complex structures (Figure S1, Table S1); analysis of target-specific scores, Vina docking scores, molecular properties and their distributions (Figure S2-S5, Table S2); comparison of generated molecules, their docking scores, and poses with previous studies (Figure S6-S7); evaluation of generation process growth (Figure S8); analysis of generated molecules in relation to active molecules and a comparison of guided generation versus REINVENT2.0 (Figure S9-S10, Table S5-S6); molecule examples and scoring model performances (Figure S11-S14); and detailed model training times and calculations (Table S3-S4).

## Acknowledgments

This work is supported by the “Pioneer” and “Leading Goose” R&D Program of Zhejiang (2023C03109), the National Natural Science Foundation of China (32171247, 21803057), the Zhejiang Provincial Natural Science Foundation of China (LQ23F020011, LR19B030001), the Central Guidance on Local Science and Technology Development Fund of Zhejiang Province (2022ZY1006), and the Westlake Education Foundation. We thank the Westlake University Supercomputer Center for computational resources and related assistance.

## MATERIALS AND METHODS

### Training of Target-specific, structure-free scoring model

A key component in the protocol is the training and utilization of target-specific, structure-free scoring models. For a given target protein, docking calculations are carried out on a curated compounds library with Vina to obtain the docking score as the labeled data. Each compound is represented by a molecular graph with atom features as nodes and bond features as edges, which is used as input to train the target-specific scoring model. Directed message passing neural network (D-MPNN)^32^ is trained with the directed edge-based message encoding by 300 dimensions to transmit information across the compounds with 3 iterations in the message passing phase, employing the chemprop library^52^. The atom hidden states are obtained by aggregating the hidden states of bonds to output the feature representations in the readout phase. The target-specific score is predicted by a feed-forward neural network by two fully connected layers with 300 dimensions. The L2 norm between Vina scores and predicted scores is used as the loss function and ReLU as the activation function. The Adam optimization algorithm is used to train our models using 50 epochs with a batch size of 50.

### Molecular generation guided by target-specific scoring model

The trained target-specific scoring models are included into the evaluation component in the molecular generative models to generate molecules that can bind favorably to the specific target. For each compound, the target-specific score *d* is transformed by a sigmoid function to be scaled between [0,1] with more negative scores closer to 1 (Fig. S14),

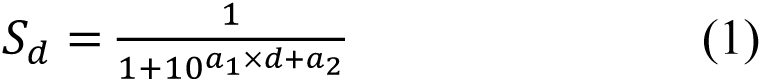

where 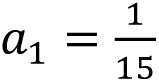, a_2_ = 1. While the molecular generation is mainly guided by *S_d_* the molecular weights and diversity is also considered in this work with following multiparameter object (MPO) score,

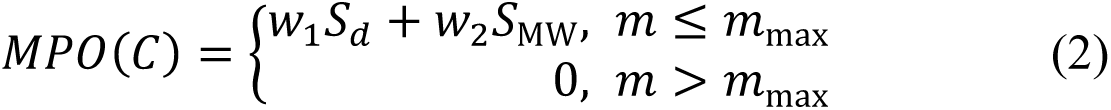

in which the weighting factors 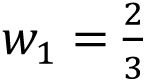 and 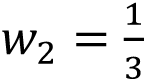. *S*_MW_ is a normalized molecular weight score that ranges close to 1 for molecules with MWs between 200 and 650 Da and for other molecules close to 0 (Fig. S14),

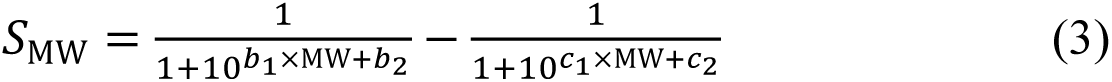

where 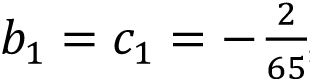, 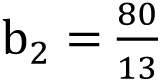 and *c*_2_ = 20.

In Eq. 2, *m* refers to the number of molecules stored in the memory that have the same Murcko scaffold with the given molecule *C*. *m*_max_ as the upper limit of *m* (set to 10 in this work) in the diversity filter controls the diversity of generated molecules.

For every epoch of molecular generation, the negative log-likelihood (NLL) is calculated for preparing loss function,

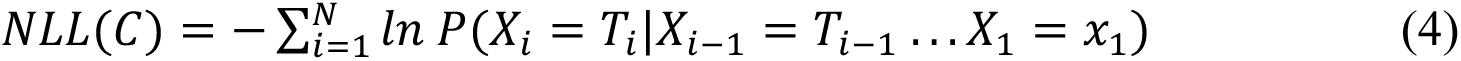

in which 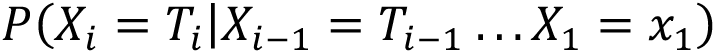 is the conditional probability of sampling a token *T_i_* at step *X_i_* given the previously sampled tokens. The ‘prior’ is a generative model which is the same as the agent at the beginning of the reinforcement learning, and the resulting MPO score is combined with the prior’s likelihood to form the augmented likelihood *NLL*(*C*)*_Augmented_*. Ultimately, the loss is calculated as the squared difference between the agent’s likelihood and the augmented likelihood.

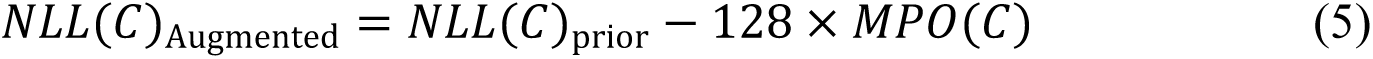

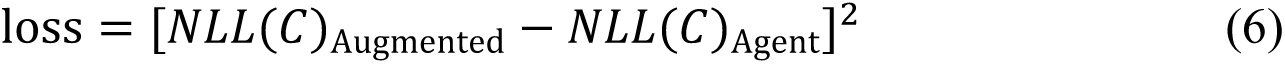

In general, the agent carries out the action to generate a sequence of tokens that build into a SMILES string. Iteratively, actions are evaluated and rewarded to generate diverse small molecules with favorable target-specific scores. The NN architecture consists of a 256-dimensional embedding layer followed by 3 LSTM layers of 512 cells and dropout layers between inner RNN layers. A liner transformation layer is used to output the vocabulary and a softmax function calculates the probabilities of every token. The batch size is 128 and all generated molecules are stored in the memory.

### Docking and compound similarity calculation

Docking calculations are performed by QVina^53^. To prepare the molecular 3D structures for docking, energy minimizations were performed with MOE^54^. MGLTools^55^ is used to translate molecules into the Vina input format PDBQT. The center of mass of the crystal ligand for each target protein is set as the center of the docking box with a varying size of 15 to 22 Å. Additional docking calculations were performed using the Vinardo docking score^46^ and DOCK6^56^. For DOCK6 calculations, default flexible docking was employed with the box size set to ensure that the largest distance between pocket grid points matches the box size used in Vina. The Tanimoto coefficient between two molecules is calculated with the ECFP4 fingerprint by RDKit. The 3D similarity is calculated by LS-align rigid body alignment^27^.

### Datasets

The commercial ChemBridge database (accessed on July 1, 2019) are used for initial docking calculations to train the target-specific scoring models. The REINVENT2.0 generative model employed in our protocol was trained using the ChEMBL library (version 22, 1.4 million compounds). Evaluations of the similarity of generated molecules were performed by comparing them with compounds in the ZINC20 dataset (accessed on July 23, 2021, 1.6 billion compounds) and the ChEMBL library (version 30, accessed on March 10, 2022, 2.1 million compounds). For each of the eight target proteins, we further collected the experimentally reported active compounds from ChEMBL library (Table S5).

### Target proteins

In our study, eight protein systems are used for evaluating the performance of the guided generative model. They include AmpC beta-lactamases (PDB ID:1L2S), dopamine receptor D4 (PDB ID:5WIU), Poly(ADP-ribose) polymerase 1 (PARP1; PDB ID:5DS3), Aldehyde dehydrogenase 1A1 (ALDH1; PDB ID:4WP7), four kinase receptors including Tyrosine-protein kinase JAK2 (PDB ID:3UGC), receptor tyrosine-protein kinase EGFR (PDBID:2RGP), Pyruvate kinase isozymes M2 (PKM2; PDB ID:5X1W) and Mitogen-activated protein kinase 1 (MAPK1; PDB ID:4XJ0). The crystal ligands binding with the target proteins are listed in Table S1.

### Enrichment factor

To compare molecular generation to docking with a given compound library, we introduce an enrichment factor from ref^5^ that provides a quantitative measurement of how generated molecules are enriched in hits compared to an enumerated library of the same size. The enrichment factor is calculated for hits with docking score better than a certain threshold *x*,

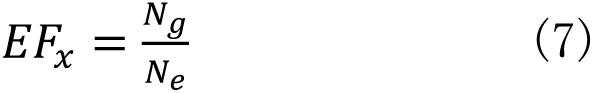

Where *N_g_* is the number of generated molecules with Vina score less than *x* and *N_e_* is the number of compounds with Vina score less than *x* in a given enumerated library.

## TOC graphics

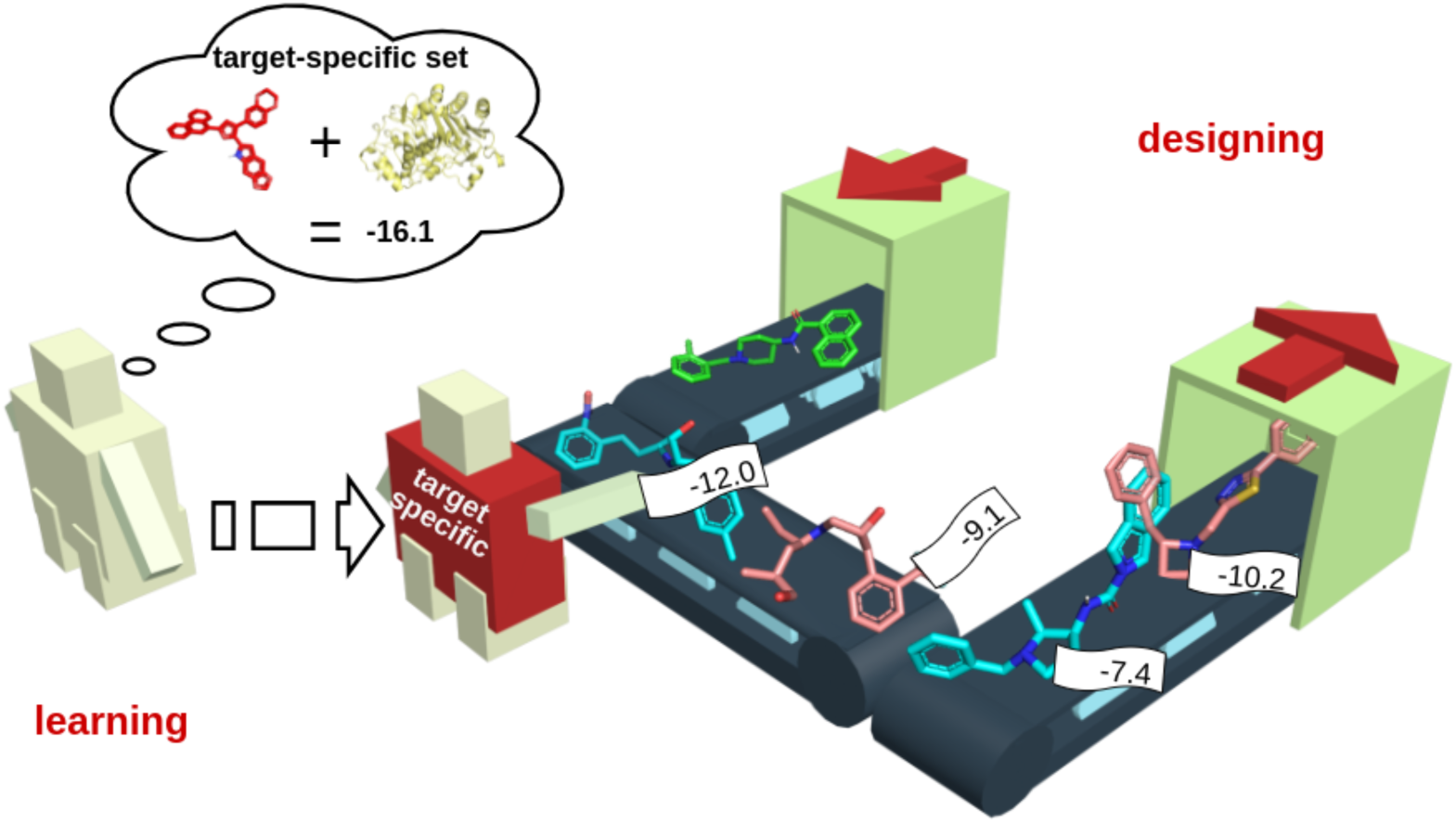

