## Supporting Information for "A Simple Way to Incorporate Target Structural Information in Molecular Generative Models"

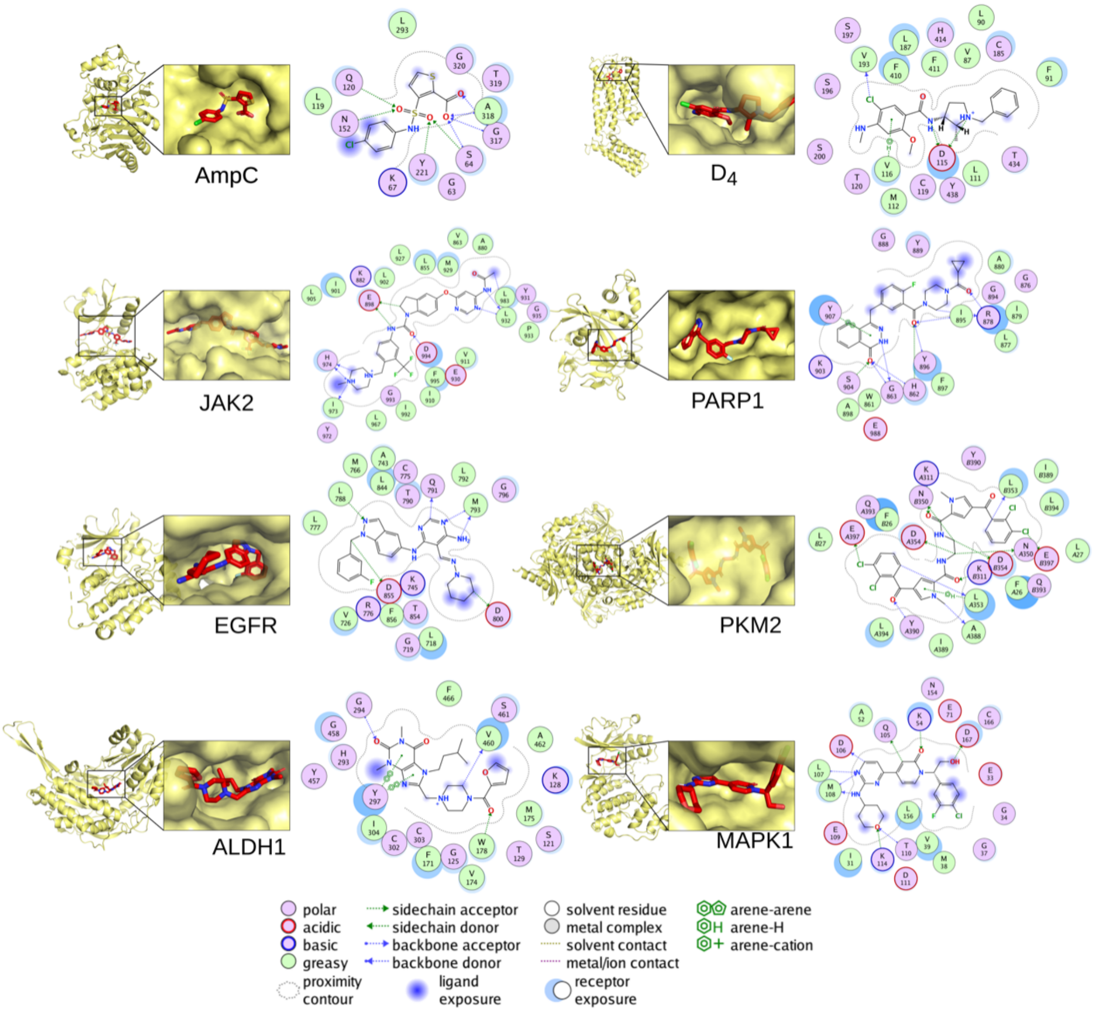


Figure S1. The experimental protein-ligand complex structures of eight target proteins in this study. Proteins and ligands were shown in yellow and red, respectively. Details on the complex structures are summarized in Table S1.

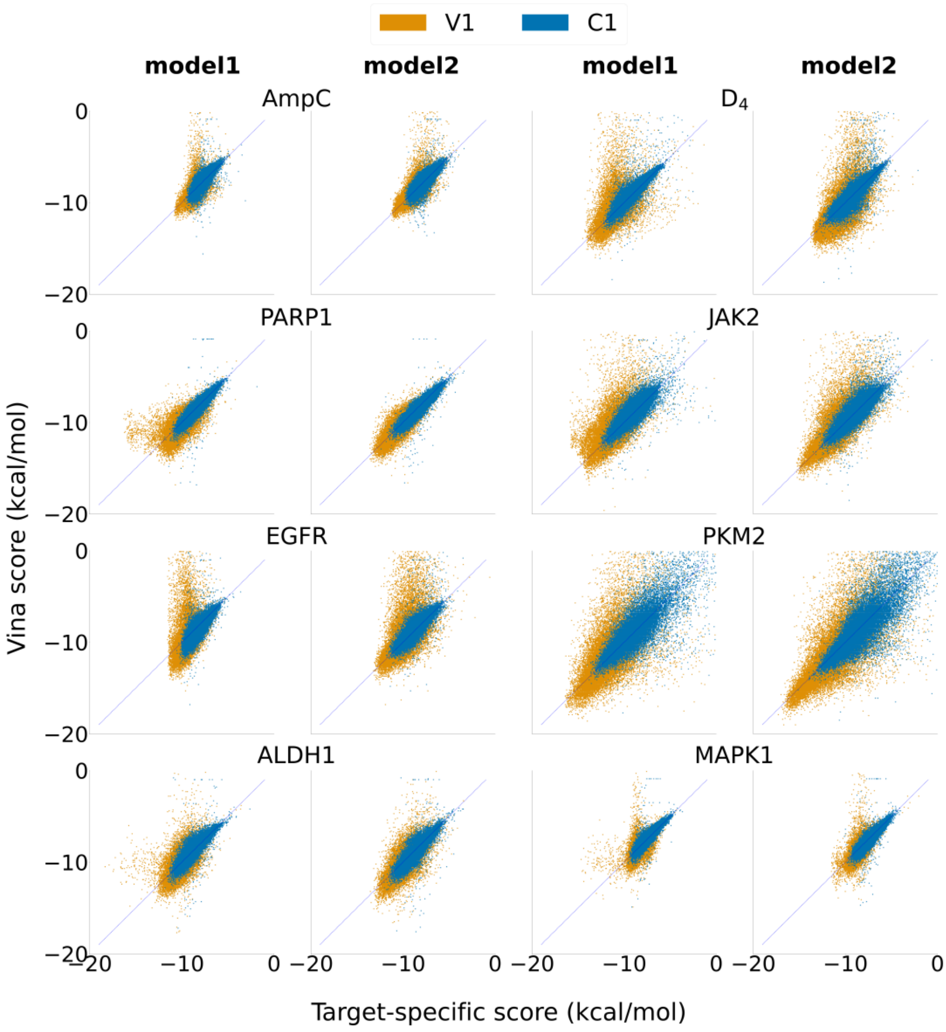


Figure S2. Comparison on the target-specific scores and Vina docking scores for eight target proteins. C1 is a subset of the ChemBridge library and V1 is a subset of generated molecules. Each set contains 50,000 compounds that are not used in training the models. Model 1 refers to target-specific scoring model trained with ChemBridge library while model 2 corresponds to training with a combined set of ChemBridge compounds and generated molecules.

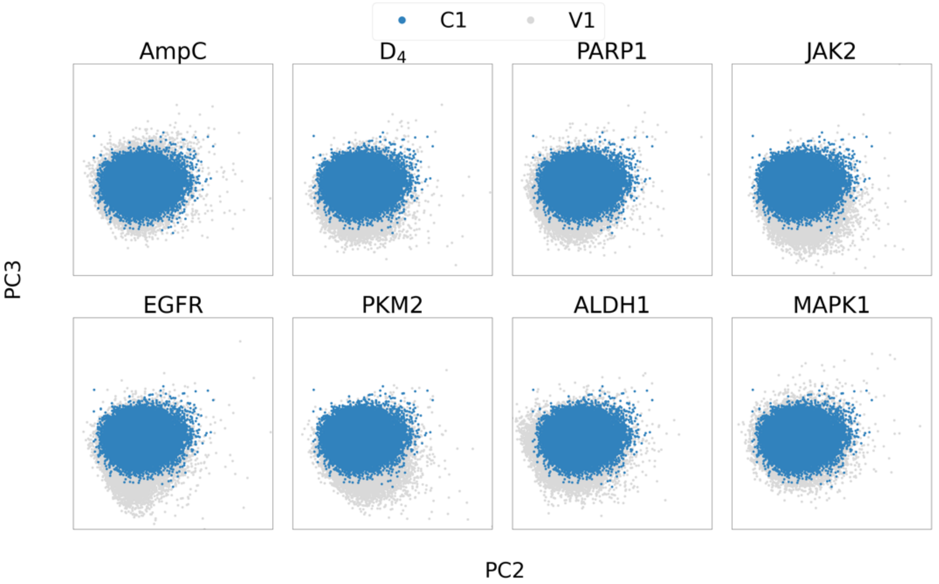


Figure S3. Comparison between the C1 (blue) and the V1 (grey) sets on the 2nd and 3rd PCs of PCA analysis using MQN descriptors.

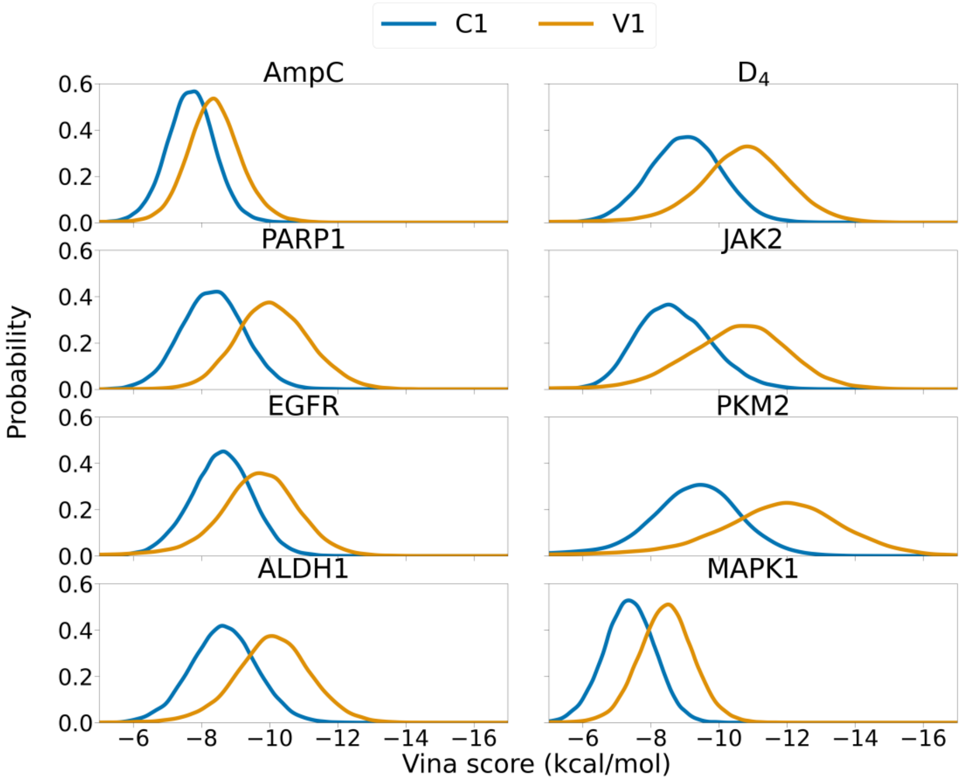


Figure S4. Comparison on the distribution of Vina docking scores of molecules in the C1 (blue) and V1 (orange) set for eight target proteins.

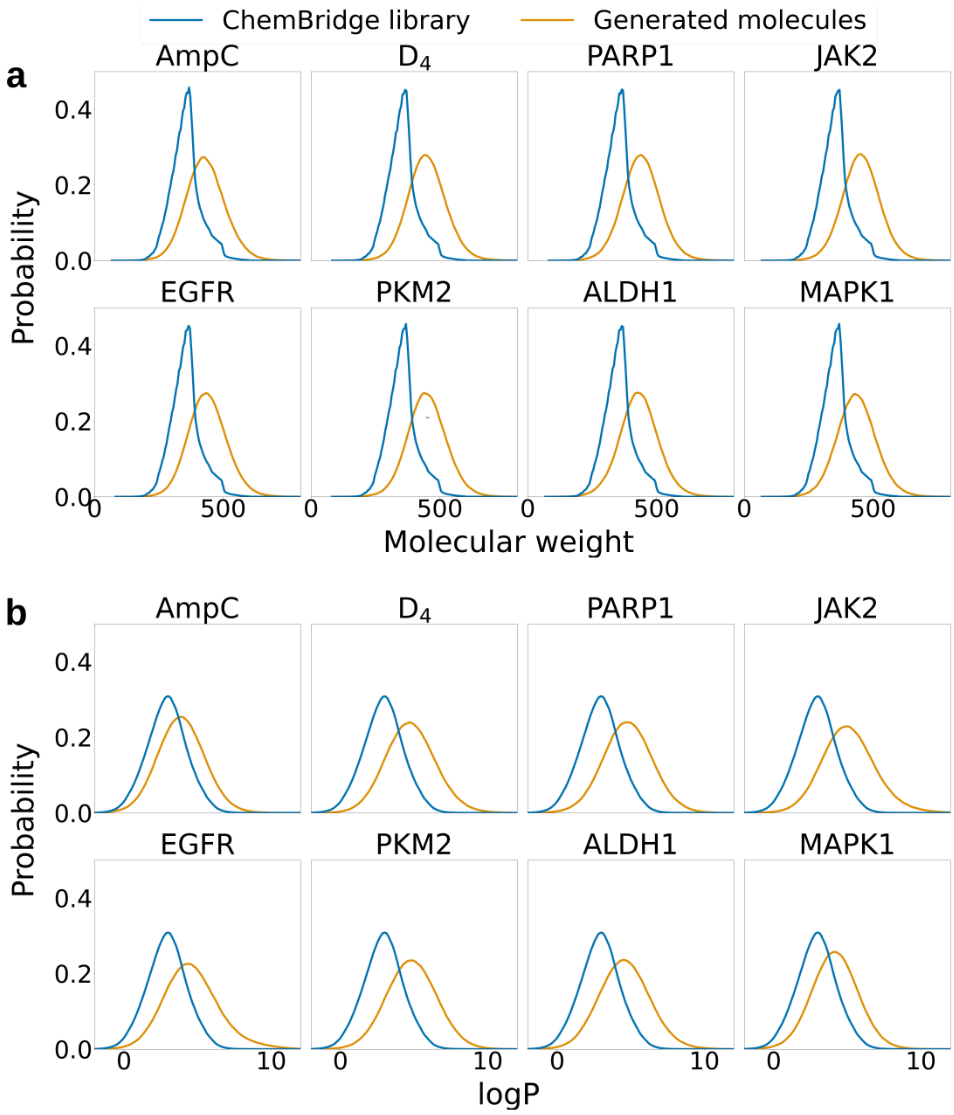


Figure S5. Comparison of (a) molecular weight and (b) logP distributions between the first generated set (orange) and ChemBridge library (blue) which both contains 743,808 molecules.

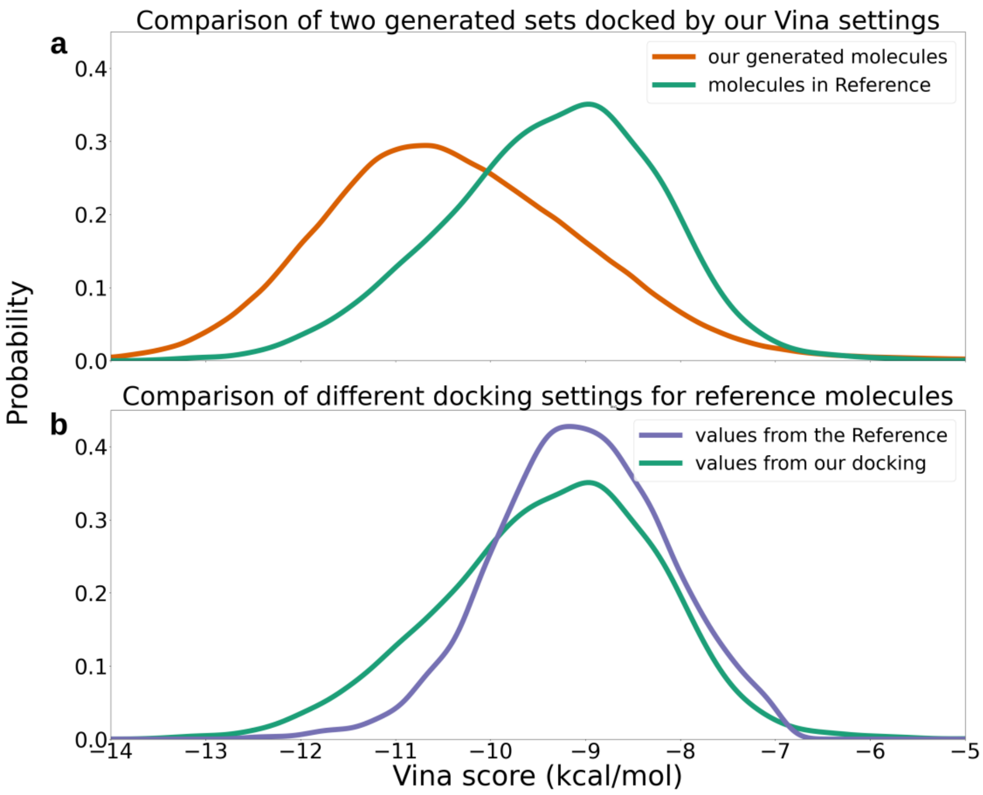


Figure S6. Comparison on the distribution of Vina docking scores for 524,282 generated molecules and the 6,106 generated molecules reported in Ref. 1. a) comparison on the distribution of Vina scores using the docking setting in this work. Green line represents the distribution for 6,106 molecules in Ref. 1 and orange line represents the distribution for generated molecules in this work. b) comparison on the distribution of Vina scores for the 6,106 molecules in Ref. 1 with different Vina docking settings. The purple line represents the values reported in Ref. 1. In general, the two docking settings lead to similar results, with slightly more favorable values with the settings used in this work. Meanwhile, the 524,282 generated molecules represent a distribution of significantly more negative Vina docking scores.

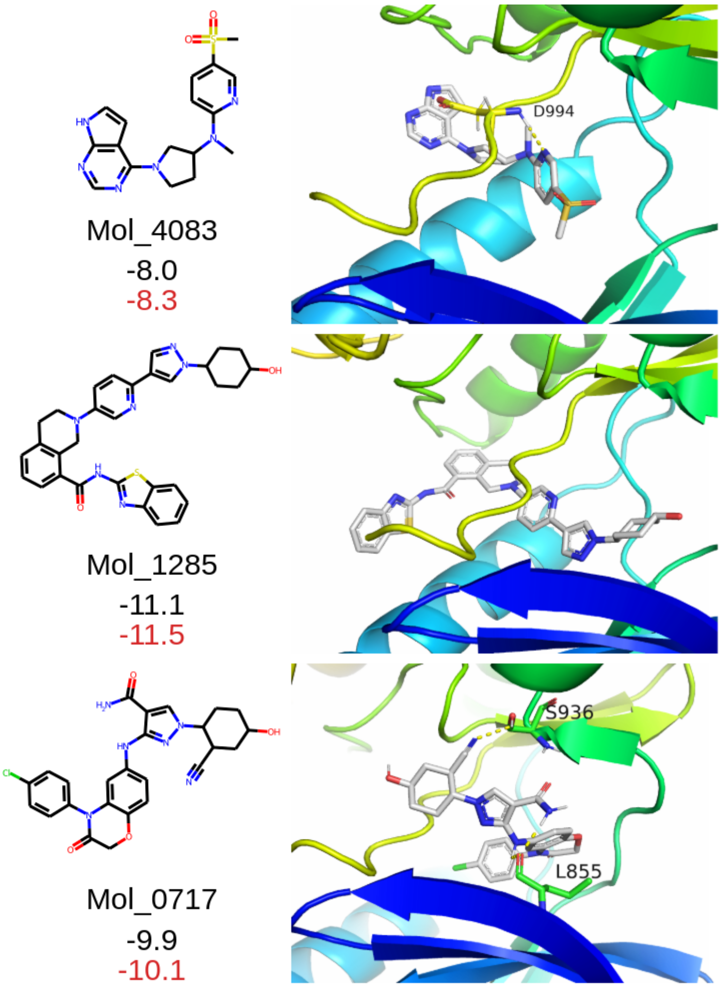


Figure S7. Comparison of docking scores and poses for example molecules reported in Ref[1] by the Vina docking setting in this work. The PDB ID of 3UGC was used in our study, compared with the 4IVA in Ref. 2. The black and red numbers (units in kcal/mol) represent docking scores for reported values and our calculations. The results were very close with slightly more favorable docking scores were obtained with the setting used in this work.

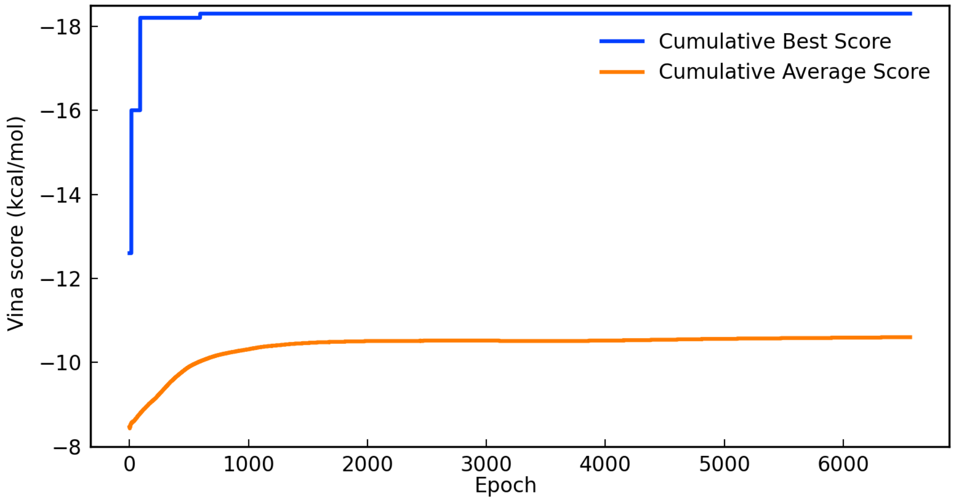


Figure S8. Cumulative average and best Vina docking scores across the generation epochs for the JAK2 target. Each epoch corresponds to the generation of 128 molecules, and a steady increase in the docking scores for generated molecules can be observed during the initial epochs.

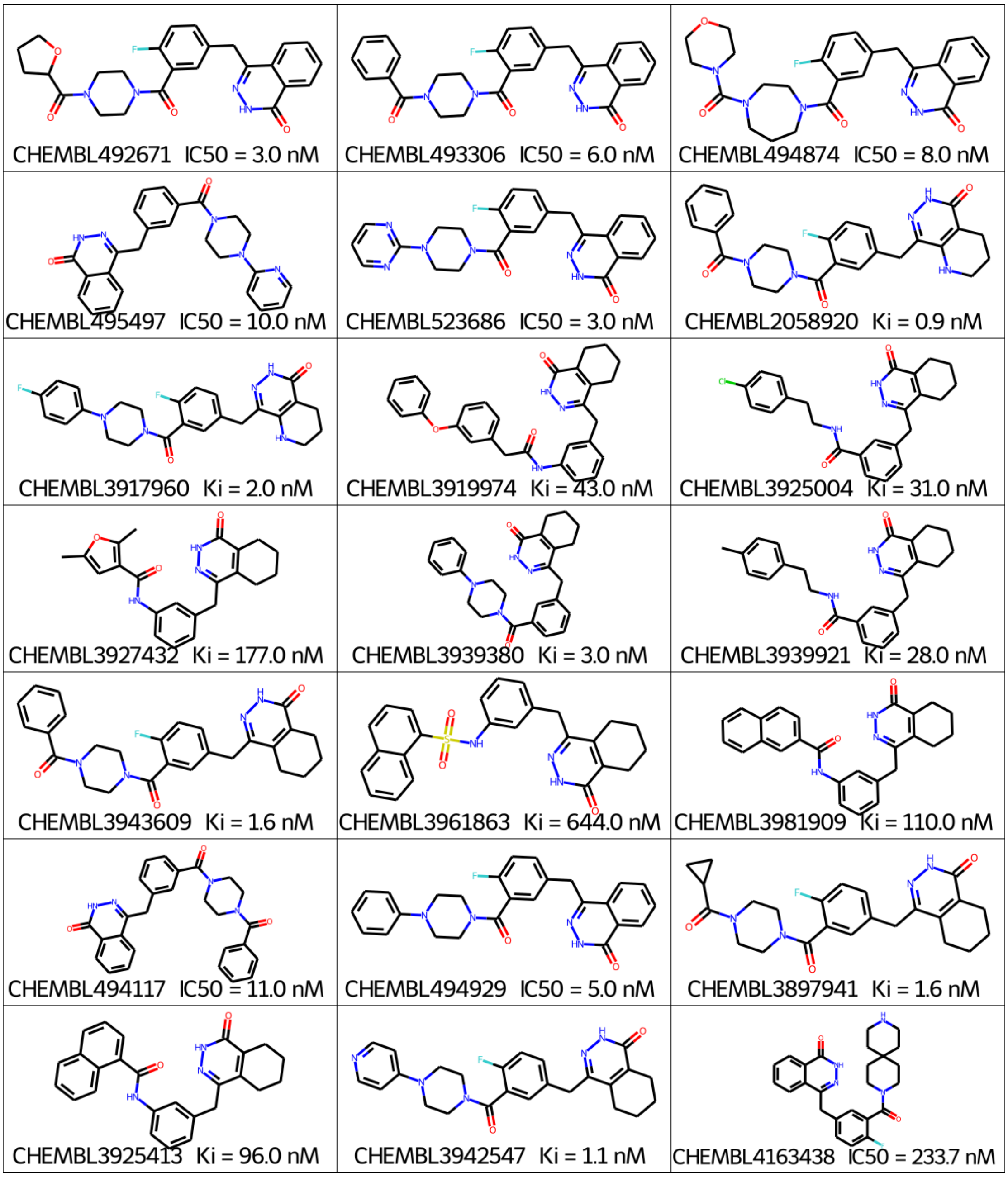


Figure S9. Twenty-one molecules generated for the specific PARP1 target that have identical structures to known inhibitors of PARP1 in the ChEMBL database.

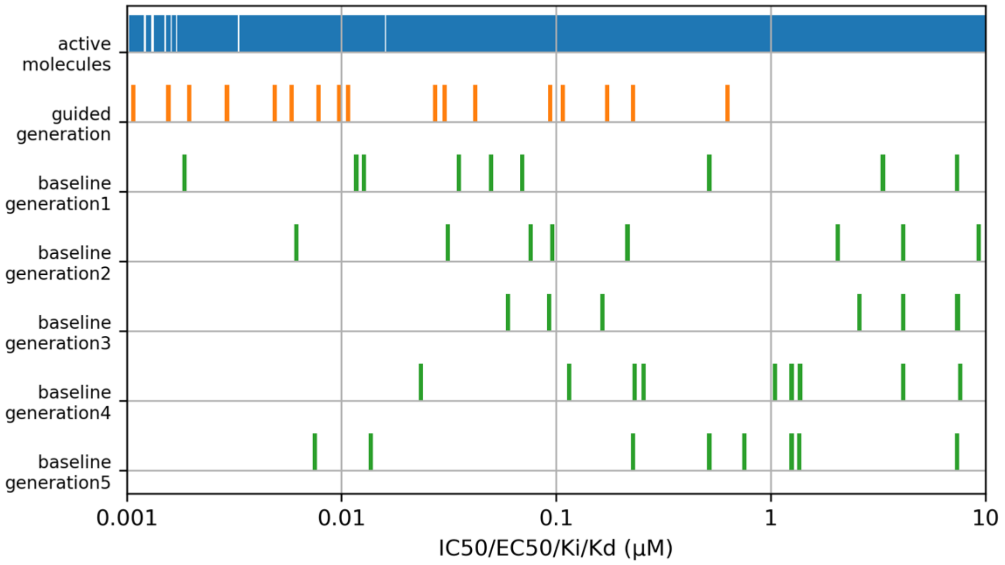


Figure S10. Comparison on the activities of recovered hits among generated molecules for the PARP1 target between target-specific guided generation (orange) and baseline REINVENT runs (green). The blue bins represent the “ground truth” affinities of the active compounds in ChEMBL for the target. The orange and green bins denote the affinities of the recovered hits by our target-specific guided generation and REINVENT, respectively. The affinities are the values of IC50, EC50, Ki or Kd collected from the ChEMBL library.

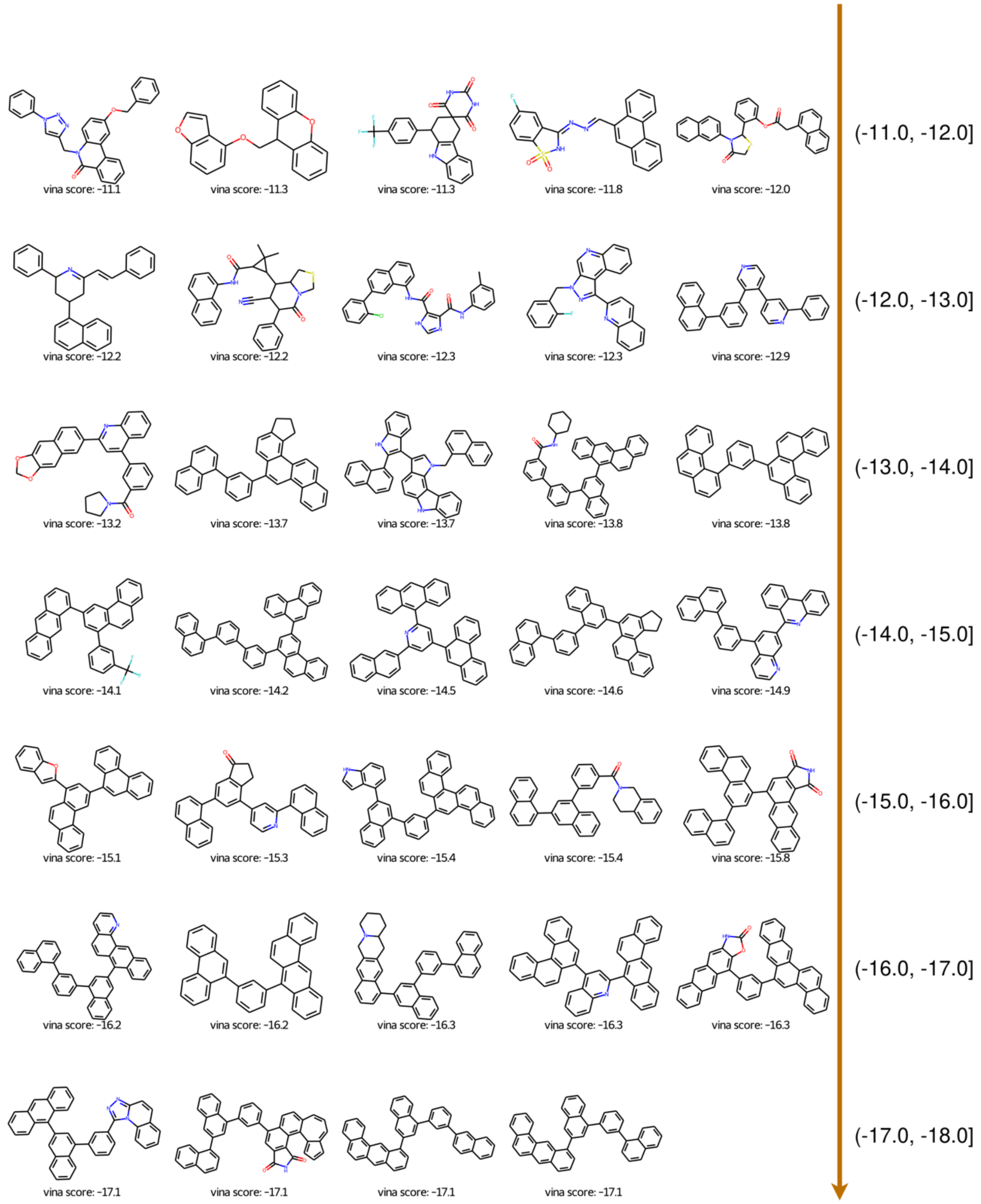

Figure S11. Example molecules with different ranges of docking scores for ALDH1.

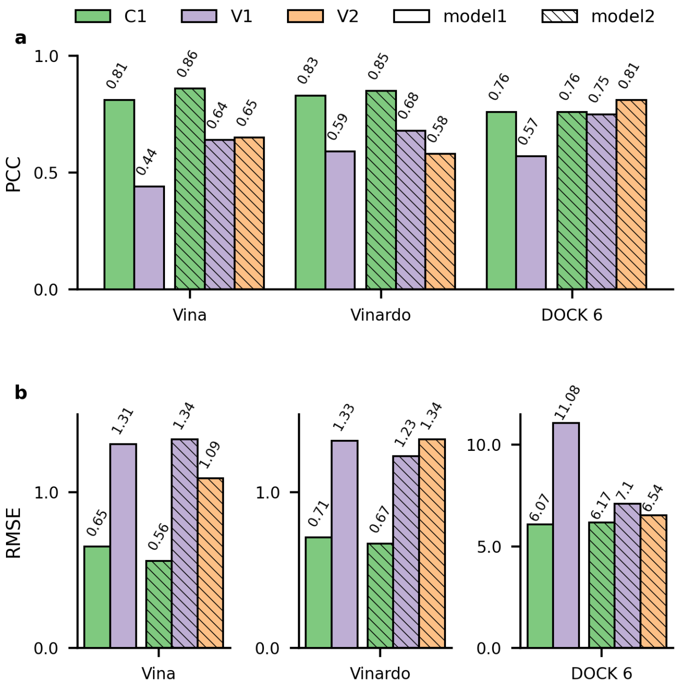


Figure S12. The PCC and RMSE of target-specific scoring models constructed using three different docking methods (Vina, Vinardo and DOCK6) for the EGFR target.

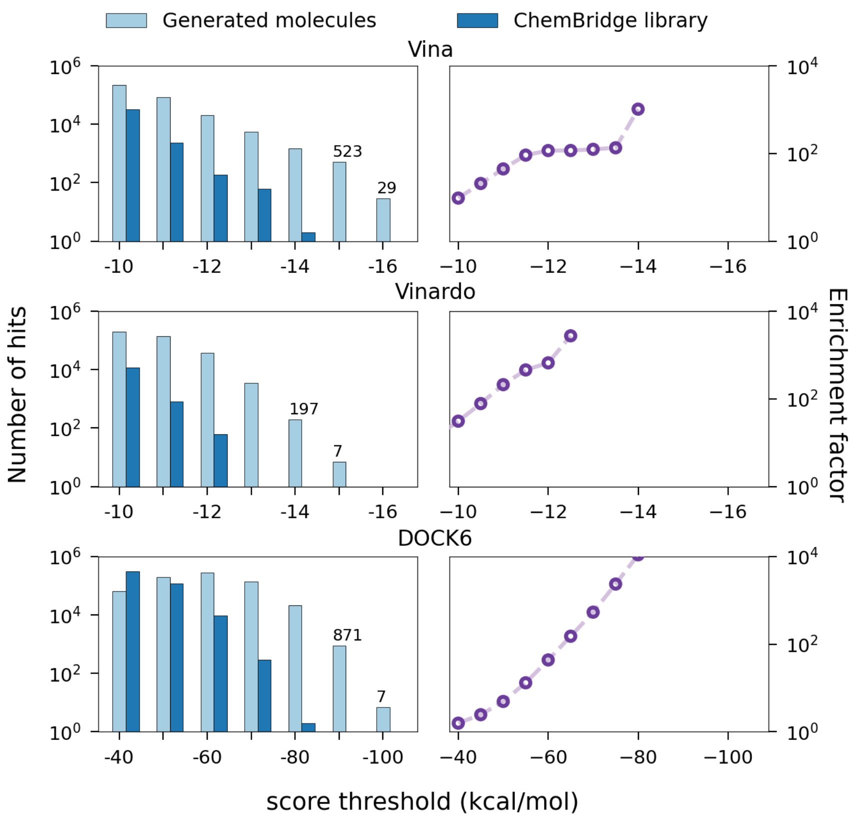


Figure S13. Comparison between generated molecules (light blue) and ChemBridge compounds (blue) with different docking methods on the EGFR target. Enrichment factors are illustrated by the purple lines. Using the docking scores of the 100th ranked ChemBridge hits as the threshold, the enrichment factors were 116, 608 and 1,116 for the target-specific molecular generation employing Vina, Vinardo, and DOCK6, respectively.

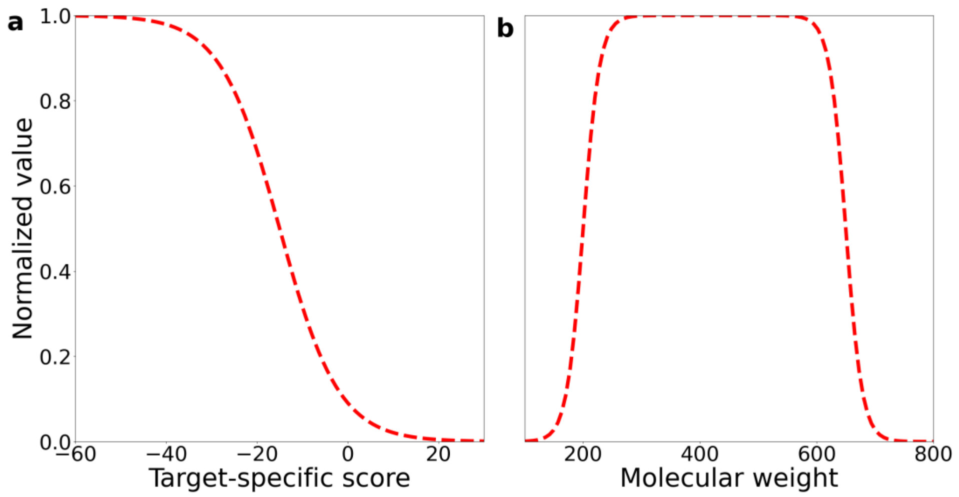


Figure S14. Illustration on the relationship between initial and transformed values of target-specific scores (a) and molecular weights (b).

Table S1. Target proteins for evaluating our protocol.

| Protein name | abbreviation | PDB ID | Resolution (Å) | Chains ID |
|---|---|---|---|---|
| AmpC beta-lactamases | AmpC | 1L2S | 1.94 | B |
| dopamine receptor D4 | $D_4$ | 5WIU | 1.96 | A |
| Poly (ADP-ribose) polymerase 1 | PARP1 | 5DS3 | 2.60 | A |
| Tyrosine-protein kinase JAK2 | JAK2 | 3UGC | 1.34 | A |
| Epidermal growth factor receptor | EGFR | 2RGP | 2.00 | A |
| Pyruvate kinase isozymes M2 | PKM2 | 5X1W | 3.00 | A, B |
| Aldehyde dehydrogenase 1A1 | ALDH1 | 4WP7 | 1.80 | A |
| Mitogen-activated protein kinase 1 | MAPK1 | 4XJ0 | 2.58 | A |

Table S2. PCC and RMSE calculations on a subset of 50,000 molecules with three additional independent samplings from the first generated set (800,000 molecules).

| Protein name | PCC | | | | RMSE (kcal/mol) | | | |
|---|---|---|---|---|---|---|---|---|
| | V1 | alternative sampling | | | V1 | alternative sampling | | |
| AmpC | 0.49 | 0.50 | 0.50 | 0.51 | 0.79 | 0.78 | 0.78 | 0.77 |
| $D_4$ | 0.55 | 0.56 | 0.56 | 0.55 | 1.27 | 1.26 | 1.27 | 1.28 |
| PARP1 | 0.69 | 0.69 | 0.69 | 0.69 | 0.89 | 0.89 | 0.88 | 0.88 |
| JAK2 | 0.64 | 0.65 | 0.64 | 0.65 | 1.37 | 1.36 | 1.36 | 1.35 |
| EGFR | 0.44 | 0.44 | 0.44 | 0.43 | 1.31 | 1.32 | 1.33 | 1.32 |
| PKM2 | 0.65 | 0.66 | 0.66 | 0.65 | 1.69 | 1.65 | 1.69 | 1.71 |
| ALDH1 | 0.61 | 0.60 | 0.62 | 0.62 | 0.97 | 0.98 | 0.95 | 0.95 |
| MAPK1 | 0.61 | 0.62 | 0.62 | 0.62 | 0.69 | 0.68 | 0.68 | 0.68 |

Table S3. Details of model training times and associated calculations taking EGFR as an example. NVIDIA A40 GPUs and Intel Xeon Gold 6150 CPUs (2.70GHz) were used.

| events | elapsed time | resources |
|---|---|---|
| Docking on ChemBridge library | 16h 20min | 900 CPU cores |
| Training D-MPNN model 1 | 1h 52min | 1 GPU, 6 CPU cores |
| Generating first 800,000 molecules | 3h 4min | 1 GPU, 6 CPU cores |
| Converting SMILES to “pdbqt” (500,000) | 38min | 500 CPU cores |
| Docking with 500,000 molecules | 15h 50min | 2,000 CPU cores |
| Training D-MPNN model 2 | 4h 57min | 1 GPU, 6 CPU cores |
| Generating second 800,000 molecules | 3h 10min | 1 GPU, 6 CPU cores |
| Total | 45h 51min | |

Table S4. Details of generated molecules with TC=1 compared to compounds in the ZINC20 library and training datasets (ChEMBL and ChemBridge), and the range of Vina docking scores of the top 20 and top 50 generated purchasable hits. The size of ZINC20, ChEMBL and ChemBridge are about 1.6 billion, 1.4 million and 0.7 million, respectively. Values in brackets indicate proportions within the training datasets.

| Target | ZINC 20 | Docking scores of top 20 hits (kcal/mol) | Docking scores of top 50 hits (kcal/mol) | ChEMBL & ZINC20 | ChemBridge & ZINC20 | ChEMBL & ChemBridge & ZINC20 |
|---|---|---|---|---|---|---|
| AmpC | 1262 | -10.6 to -11.1 | -10.4 to -11.1 | 93 (0.006%) | 46 (0.006%) | 13 |
| D4 | 275 | -13.2 to -13.7 | -12.9 to -13.7 | 14 (0.001%) | 15 (0.002%) | 2 |
| PARP1 | 124 | -12.2 to -13.2 | -12.0 to -13.2 | 4 (0.0003%) | 4 (0.0005%) | 2 |
| JAK2 | 130 | -13.2 to -14.7 | -12.9 to -14.7 | 9 (0.0006%) | 8 (0.001%) | 2 |
| EGFR | 286 | -12.1 to -13.0 | -11.9 to -13.0 | 19 (0.001%) | 18 (0.002%) | 3 |
| PKM2 | 127 | -14.7 to -15.8 | -14.3 to -15.8 | 3 (0.0002%) | 5 (0.0007%) | 1 |
| ALDH1 | 318 | -13.0 to -14.2 | -12.5 to -14.2 | 27 (0.002%) | 19 (0.003%) | 4 |
| MAPK1 | 324 | -10.3 to -12.7 | -10.0 to -12.7 | 32 (0.002%) | 8 (0.001%) | 1 |

Table S5. Collection of active molecules and numbers of generated molecules similar to active molecules for the eight target proteins. The values in brackets indicate the proportion in the training datasets which include about 1.4 million and 0.7 million for training REINVENT and D-MPNN models, respectively. Affinity refers to the values of IC50, EC50, Ki or Kd.

| Protein | No. of collected actives with affnity<10μM | No. of generated molecules by Tanimoto similarity>0.7 with actives |
|---|---|---|
| AmpC | 160 | 0 |
| $D_4$ | 2431 | 23 |
| PARP1 | 3112 | 279 |
| JAK2 | 5401 | 1 |
| EGFR | 6205 | 4 |
| PKM2 | 2134 | 7 |
| ALDH1 | 13421 | 45 |
| MAPK1 | 7382 | 26 |

Table S6. Comparison on recovered hits of generated molecules with actives for the PARP1 target between target-specific guided generation and five random runs of REINVENT.

| Dataset | Number of recovered hits to the PARP1 | Number of generated molecules that were found in the ChEMBL database | percentage |
|---|---|---|---|
| guided generation | 21 | 53 | 39.62% |
| baseline generation 1 | 9 | 4450 | 0.20% |
| baseline generation 2 | 10 | 4451 | 0.22% |
| baseline generation 3 | 9 | 4641 | 0.19% |
| baseline generation 4 | 9 | 4333 | 0.21% |
| baseline generation 5 | 9 | 4605 | 0.20% |
